# Multiscale model of the different modes of cancer cell invasion

**DOI:** 10.1101/2022.10.07.511296

**Authors:** Marco Ruscone, Arnau Montagud, Philippe Chavrier, Olivier Destaing, Isabelle Bonnet, Andrei Zinovyev, Emmanuel Barillot, Vincent Noël, Laurence Calzone

**Affiliations:** Institut Curie, Université PSL, F-75005, Paris, France; INSERM, U900, F-75005, Paris, France; Mines ParisTech, Université PSL, F-75005, Paris, France; Sorbonne university, Paris; Barcelona Supercomputing Center (BSC), Barcelona, Spain; Institut Curie, PSL Research University, CNRS, UMR 144, Paris, France; Institute for Advanced Biosciences, Centre de Recherche Université Grenoble Alpes, Inserm U 1209, CNRS UMR 5309, France; Institut Curie, Université PSL, Sorbonne Université, CNRS UMR168, Laboratoire Physico Chimie Curie, Paris, France

**Keywords:** Multiscale, Cancer, Agent-based model, Invasion

## Abstract

Mathematical models of biological processes implicated in cancer are built using the knowledge of complex networks of signaling pathways, describing the molecular regulations inside different cell types, such as tumor cells, immune and other stromal cells. If these models mainly focus on intracellular information, they often omit a description of the spatial organization among cells and their interactions, and with the tumoral microenvironment. We present here a model of tumor cell invasion simulated with PhysiBoSS, a multiscale framework which combines agent-based modeling and continuous time Markov processes applied on Boolean network models. With this model, we aim to study the different modes of cell migration by considering both spatial information obtained from the agent-based simulation and intracellular regulation obtained from the Boolean model. Our multiscale model integrates the impact of gene mutations with the perturbation of the environmental conditions and allows the visualization of the results with 2D and 3D representations. The model successfully reproduces single and collective migration processes and is validated on published experiments on cell invasion. *In silico* experiments are suggested to search for possible targets that can block the more invasive tumoral phenotypes.

## 1. Introduction

Cancer is the top cause of disease burden in the world according to the World Health Organization (WHO) (1). The severity of cancer is greatly increased by the invasion of cancerous cells into its surrounding environment, through a metastatic process that may end up in the generation of a secondary cancer seeding (2, 3). Cell invasion is a complex multiscale process that requires the coordinated action of entities at the cells’ molecular, cellular and population level. In addition, cell invasion depends on the interplay between cells and the extracellular environment (4): the presence of proteins affect the density of the extracellular matrix (ECM). Likewise, the ECM can constrain cellular invasion (5, 6): the ECM stiffness and the availability of oxygen or nutrients impact intracellular mechanisms, and ECM acts as a repository for a variety of growth factors and matricellular proteins released upon ECM modification (7). This complex dynamic system causes a wide array of different invasion behaviors that have been described in different cancers (2, 8, 9).

Three main invasion modes have been categorized (10, 11), where cells move either individually, collectively or in streams of cells. Although these terms are arguably arbitrary and their description can be incomplete, notably at the molecular level, they are useful as they simplify and categorize the literature, and they facilitate the study of the molecular mechanisms underlying each mode (10). Single-cell or individual migration is the invasion mode where cell-cell junctions have been lost and cells are free to degrade and roam the ECM (11). Depending on the tissue, this single-cell mode can be further described as amoeboid, where cells have low adhesion force and spherical shapes, or as mesenchymal, where cells have cytoskeletal protrusions, adhesion capabilities and a spindle shape, with elongated morphologies and proteolytic activity toward ECM substrates. Collective migration is the invasion mode where cell-cell adhesions are retained and cells invade as multicellular groups, requiring a certain coordination between cell-cell adhesion and migration (12). This mode has been identified as more aggressive and with higher metastatic potential, especially in circulating tumor cells and breast cancer (13, 14). Trail or multicellular streaming migration is a special case of the single-cell invasion mode where individual cells move one after the other using the same path within the tissue (10). This can be caused by a given chemoattractant or due to local ECM heterogeneities that allows for a path of less friction for cells. Independently of the invasion mode, the first step of cellular invasion is the detachment of cells from its neighboring tumor tissue. The epithelial-to-mesenchymal transition (EMT) is a molecular program that triggers tumor cell invasion in response to environmental signals (15) by detaching cells from its neighbors (11), promoting cell-matrix adhesion and the formation of protrusion at the cell membrane (16, 17) and by secreting matrix metalloproteases (MMPs) that degrade the ECM (18). EMT allows the cell to switch to a more motile phenotype, losing adhesion to neighboring cells and promoting invasiveness at the single cell level.

To study the invasion process that spans many time and spatial scales, multiscale models such as agent-based modeling are ideally suited as they consider agents as surrogate of cells that move, divide, and die, which facilitates the specific description of both intracellular, secretion and microenvironment behaviors (19). Agent-based modeling has been successfully applied to cancer studies (20) and with the increased capacity provided by the high-performance computing, it is now efficient enough to scale to the tissue level (21). Some agent-based studies have focused on describing the effect of the pressure and the interactions between the agents and the ECM (22) without considering intracellular pathways. Other works have combined the agents with partial and ordinary differential equations to describe the gene regulatory networks of intracellular regulations (23). However, thys type of dynamical models only cover a small part of the biological mechanisms and often require many parameters that need to be fitted with experimentsOne way to cope with this limitation is to describe the intracellular regulations with the Boolean formalism, which although providing a less detailed description, requires no or very few parameters (24). In this study, we use PhysiBoSS (25, 26) a software that combines PhysiCell (27), an agent-based modeling framework that integrates interactions between cells, cell types and with the microenvironment, and MaBoSS (28, 29), a tool that relies on stochastic simulations of Boolean intracellular signaling models. PhysiBoSS allows for the combined genetic and environmental perturbations of tumors and inspects their effect at the population level, enabling the study of drug treatments and cellular heterogeneity and their effect on cancer phenotypes. PhysiBoSS is used to describe a novel model of tumor growth that combines a description of the signaling pathways that trigger events leading to an invasive phenotype, focusing on both single-cell and collective features. By varying the parameters of the agent-based model and various intracellular components, the model reproduces the different modes of invasion and provides a tool for the suggestion of potential drug treatments and genetic perturbations that could block or perturb specific invasion modes.

## 2. Methods

### 2.1 PhysiBoSS - a multiscale framework combining Boolean and agent-based models

The model of the different modes of invasion links two formalisms, providing two levels of description: one at the level of the individual cell and one at the level of cell population. PhysiBoSS allows this multiscale description combining simulation of Boolean models using continuous time Markov process (with MaBoSS), and agent-based modeling of physico-chemical cell-cell and cell-environment interactions (with PhysiCell). The multiscale model presented here combines an intracellular gene regulatory network with several signaling pathways and generic cell-level parameters. The intracellular model represents interactions often deregulated in cancer and is a refinement and extension of already published models of the invasion mechanisms (30, 31). There is a feedback of variables between the Boolean model and the agent-based model. The intracellular model receives diverse stimuli from the environment by activating a membrane receptor (input) that triggers signaling pathways and molecular interactions, ultimately leading to different cell responses through MaBoSS simulations. These cell fates are represented by phenotypes that, among others, include cell division (by activating the M-type cyclins), cell death (by cleaving the effector caspase 3) and EMT (see supplementary file). Some components can be secreted by the cell, released in the cell environment, which can activate or inactivate the neighboring cells. These cell fates, in turn, affect different variables and parameters of the agent-based model and of neighboring cells. For instance, the release of MMPs will degrade the ECM, and subsequently free some proteins that will activate the receptors of neighboring cells or the intracellular model can determine the loss of adherence with other neighboring cells and the detachment from the primary tumor.

### 2.2 The construction of the intracellular model

The intracellular model builds upon two published models, one of the early steps of metastasis (31) and another of the EMT process (30). The initial model of Cohen and colleagues was built with two inputs: the *ECMenv*, which monitored the status of the extracellular matrix, and *DNA_damage*, which considered DNA alterations that trigger death signals. Four additional inputs were added to account for the presence of *Oxygen*, growth factors (as *GF*), *TGFbeta* and the contact with other neighboring cells (as *Neigh*)(Fig. 1). The phenotypes, or outputs of the model include *CellCycleArrest, Apoptosis, EMT, ECM_adh* (for cell adhesion), *ECM_degrad* (for cell degradation), *Cell_growth* (for the dynamics of the tumor growth) and *Cell_freeze* (for cell motility ability). New genes and pathways include mechanisms around p63 (32) and SRC (33, 34). Genes from the Hippo pathway and RhoGTPases, such as YAP1 (35), FAK and RAC1 (36) were also inserted to link external signals (i.e., cell–cell contact, stiffness of the extracellular matrix, and stress signals) and intracellular regulation. Currently, the model does not include a mesenchymal to epithelial transition (MET) (37, 38), and a cell in a mesenchymal cell cannot revert its state. The resulting network encompasses 45 nodes, with 6 input nodes, representing the possible interactions of an individual cell with external elements, and 8 output nodes or read-outs describing the possible observed phenotypes (Fig. 1). The model is also provided in SIF format to facilitate the network visualization with Cytoscape (34) (see supplementary material).

**Fig. 1.**
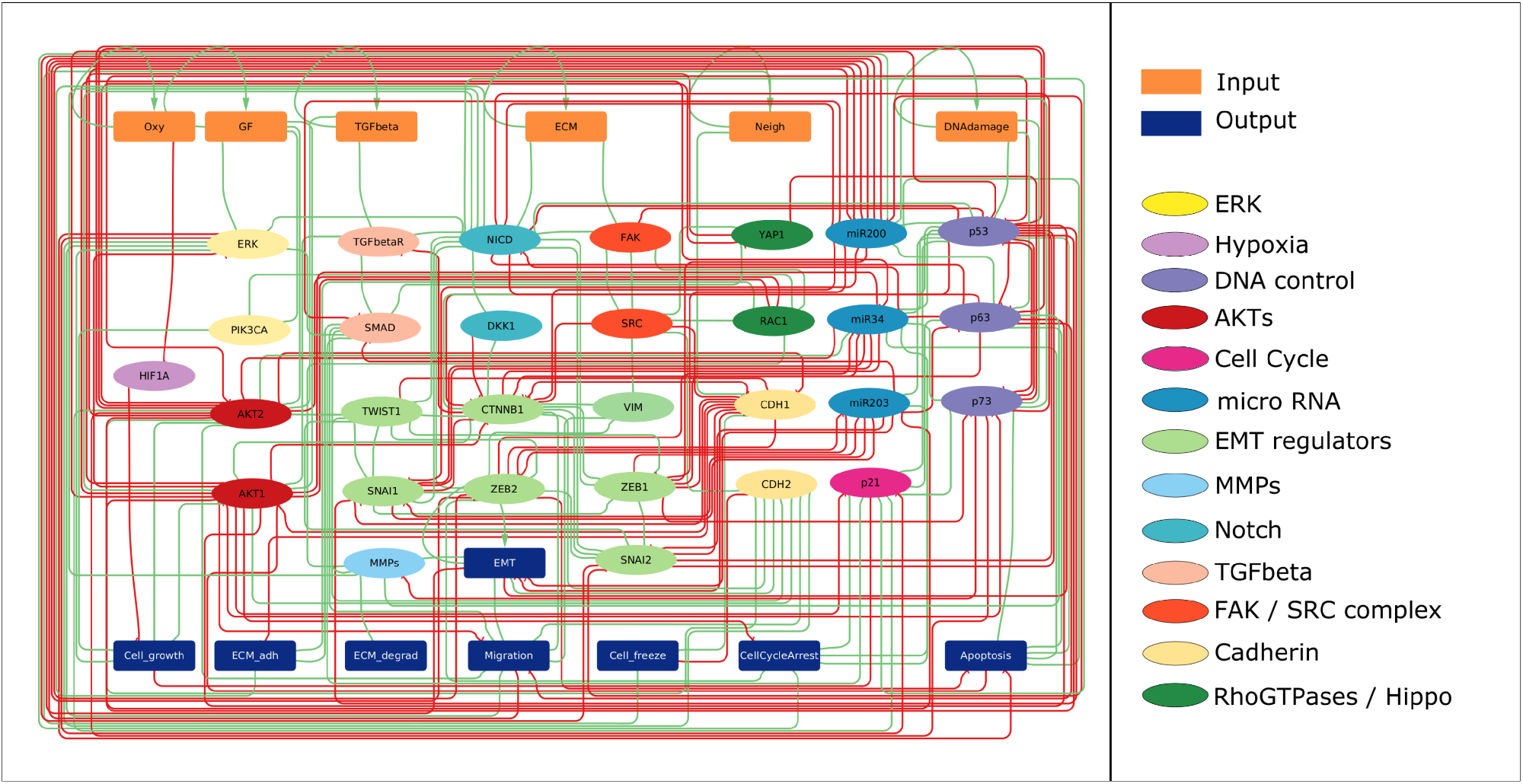
Influence network of the intracellular model. Rectangular orange nodes represent inputs, rectangular blue nodes represent outputs (or phenotypic read-outs). Ellipsoid nodes are genes or proteins that participate in the signaling pathways listed in the legend

### 2.3 Initial conditions for the simulation of the intracellular model

The initial conditions of the model were set to represent a given cell’s position in the tumor: in the center of the tumor with cells surrounding it (*Oxygen* and growth factors are considered ON), and at the periphery of the tumor in direct contact with the ECM (*Oxygen*, growth factors, *ECM* and *TGFbeta* ON with or without *DNAdamage*) (Fig. S2). It is assumed that on the surface of the tumor, cells are more likely to switch from an epithelial to a mesenchymal state, while in the center, they form tight cell junctions preventing movement. For the inner tumor cell conditions, the simulation with MaBoSS shows a high proportion of cells with *Cell_freeze* and *Cell_growth* phenotypes, corresponding to an increase in the tumor mass without signs of invasion (Fig. S2A). Instead, activation of *DNA_damage* leads to the activation of the apoptotic pathway (see Supplementary Material). The pressure due to cell crowding activates the *DNA_damage* node, which leads to the secretion of MMPs and the remodeling and degradation of ECM (39). In the absence of DNA damage, a high probability of cells with the *EMT/Migration/ECM_adh* phenotypes is observed, corresponding to a high invasive potential. This is also true in the absence of oxygen and growth factor. The presence of DNA damage inhibits *EMT* while promoting secretion of MMPs (Fig. S2C). The combination of the absence of GF and DNA damage leads to an apoptotic state (see Supplementary Material).

Different combinations of initial states can be simulated using our dedicated Jupyter notebook (supplementary material) and in the GitHub repository https://github.com/sysbio-curie/Invasion_model_PhysiBoSS.

### 2.4 Set-up of the agent-based model for the integration of the intracellular model

At the beginning of the simulation, the tumor is represented as a spheroid composed of cancer cells in an epithelial state, surrounded by the ECM. The oxygen can freely diffuse into the microenvironment towards the center of the spheroid, while TGFbeta is encapsulated into the ECM. In the model, oxygen is needed for both cellular respiration and as a triggering signal for the cells’ motility. Oxygen can freely diffuse in the environment and is uptaken by the cells. When the internalized value of the oxygen is above a threshold (*oxygen_necrotic_threshold*), the *Oxygen* node of the intracellular model is set to ON. The extracellular matrix (ECM) is defined as a static substrate, i.e., not characterized by a diffusion coefficient and without any decay that prevents the cells from moving across it. It is only when the cells have acquired the appropriate phenotype (*ECM_degrad* and *Migration* active) that they can trigger the invasion process. Secondly, the ECM contains a certain amount of TGFbeta, which is released when the cell degrades the ECM beyond a certain threshold. We fitted this threshold to reproduce experimental data (section 3). It is possible to further explore the model behaviors using different threshold values, however this parameter does not seem to affect the different modes of invasion. When ECM is degraded, TGFbeta becomes accessible to the neighboring cells that initiate its uptake. The presence of oxygen and TGFbeta in the microenvironment, if above a certain threshold, can trigger the activation of the corresponding input node (*TGFbeta, Oxy*) in the intracellular model. The ECM input node is triggered when a cell has more contact with the ECM than a user-defined threshold. This value is stored into the variable *ecm_contact* and is based on the amount of overlap between the cell and the voxel. The intracellular node *Neigh* is the readout for cell-cell contact. In order to quantify cellular contact, the distance between neighboring cells is stored into a variable of the environment *cell_contact* and corresponds to the percentage of their overlapping cell radii. This value is compared to *cell_cell_contact_threshold* thresholds to set *Neigh*’s value. To simulate DNA damage, we consider the physical stress that the cells at the border of the tumor have when they are pushed against the ECM barrier, suffering high nuclear pressure. We represent this pressure as an overlap (distance) between the nucleus radius of the agents and the voxel containing the ECM. When this overlap reaches a given threshold (*DNA_damage_threshold* = 0.8), it triggers the *DNA_damage* node.

Similarly, the outputs of the model are usually activated by a combination of active nodes and are connected to functions inside each agent. The *Apoptosis* node triggers the apoptotic death model of PhysiCell. *Migration* changes the direction of the motility of the cells, from a random walk to a chemotaxis movement towards the highest concentration of nutrients (represented by oxygen). As *Migration* is activated, the internal variable of the cell *pmotility* increases. This variable changes the length of motility so that the more a cell migrates, the more the displacement of this cell increases at each new step in the simulation. *EMT* is a read-out in the model, considered as a necessary early step for cell invasion that modifies some mechanical aspects of the agent: once activated, it impacts the *padhesion* variable that indicates the percentage of cell adhesion with neighboring cells. *EMT* value also determines whether a cell forms a cluster with neighboring cells or not, a phenomenon that is characterized by a hook-like adhesion.

The formed cluster is disbanded if: 1) *padhesion* is below a certain threshold, simulating the lack of cell junctions by the E-cadherins, or 2) an external mechanical force causes the separation of the cluster. The node *ECM_adh*, that accounts for the junction between the cell and the ECM activates a function that increases the amount of integrins in the agent (corresponding to the parameter *integrins* in the model). The node *ECM_degrad* monitors the secretion of the metalloproteases (MMPs), enzymes that allow the cells to degrade the ECM and hence start invasion. In the model ECM_degrad triggers the uptake rate of ECM substrates based on the amount of integrins displayed by a cell, reducing the density of the ECM value in the target voxel and facilitating the cell’s movement through it.

### 2.5 Impact of the parameter values on the different modes of invasion

The different modes of invasion depend on the values of a set of parameters (see supplementary file), which correspond to biophysical mechanisms difficult to infer from experiments. A set of parameter values is proposed in order to simulate the different invasion modes, qualitatively reproducing the experimental images (section 3). To study the impact of the main parameter values, we performed a sensitivity analysis on a subset of 7 parameters, that cause the simulation to switch from one mode of invasion to another (Fig. 4). For each parameter, we select a range of values (Table 1) and run 50 replicates for each. By varying their value, we are able to modulate the presence of single cells and cells migrating in clusters (Supplementary file). The 7 parameters are listed in Table 1.

**Table 1.** List of the 7 parameters studied in the sensitivity analysis. For each parameter, we focused on a range of values, picked several values in it, ran 50 simulations for each parameter set and ended up selecting the number in bold (see Supplementary material).

| Parameters | Description | Range |
| --- | --- | --- |
| *cell_ecm_repulsion* | regulates the amount of repulsion between cell and ECM | 5 < **15** < 40 |
| *epith_cell_attach_threshold* | changes the activation threshold needed to attach cells in cluster with cell junction | 0.001 < **0.05** < 1 |
| *mes_cell_detach_threshold* | change the activation threshold needed to detach cells in cluster with cell junction | 0.001 < **0.03** < 1 |
| *cell_cell_contact_threshold* | changes the activation threshold of the value cell_contact needed to trigger the Neigh node | 0.01 < **0.3** < 3.5 |
| *cell_ecm_contact_threshold* | changes the activation threshold of the value ecm_contact needed to trigger ECM node | 0.001 < **0.05** < 1 |
| *migration_bias* | changes the value of migration bias for cells with Migration node active | 0.5 < **0.85** < 1 |
| *migration_speed* | changes the value of migration speed for cells with Migration node active | 0.3 < **0.7** < 1 |

### 2.6 Single and collective cell migration

Single cell migration occurs when a cell has lost the cell junctions with the surrounding cells and starts the invasion process on its own. Collective migration in cancer can happen when one of two dynamical features appear: (1) a group of epithelial cells go through EMT and start degrading a specific area of the ECM without developing cell junctions before migrating along extracellular boundaries (i.e., fibers, basal membrane, etc. that cannot be degraded) in an uncoordinated collective movement (40); or (2) a group of mesenchymal cells surrounding a small cluster of epithelial cells detach themselves from the main body of the tumor, thus, when epithelial cells have cell junctions with other mesenchymal cells which push them through the ECM. The parameter *migration_bias* governs the stochasticity of the movement of mesenchymal cells. The higher its value, the less stochastic the movement is. From the sensitivity analysis results, a significant increase in the number of single cells is observed, with values of *migration_bias* higher than 0.75: with less noise in their movement, the cells reduce their probability to meet others and to form clusters. This behavior change for values of *migration_bias* between 0.3 and 0.75. In this case, we observed that the percentage of cells in clusters is higher than single cells (around 60%, Fig. S10). A similar result is shown for the *migration_speed* parameter. Increasing the speed of mesenchymal cells leads to an increase of migrating single cells. The number of single cells grows linearly from a *migration_speed* value of 0.3 to 0.8 (Fig. S11). In biological terms, we can conclude that the more motile a cell is, the less likely it will form a cluster. The range over which to vary the threshold value for *cell_ecm_contact_threshold* and *cell_cell_contact_threshold* was calculated based on the maximum compression reached between cell/cell and cell/ECM. The analysis on the *cell_ecm_contact_threshold*, which represents the ability of the cell to detect ECM, did not show an impact on the migration mode but on the amount of invasion. For higher values (>0.5), we observe a drop of the migrating cells, whereas for lower values, the ratio between single cells and cells in clusters remains constant. For *cell_cell_contact_threshold*, the model proved to be very robust, with no significant impact on the amount or mode of invasion. The parameters *epith_cell_attach_threshold* and *mes_cell_detach_threshold* monitor the process of junction formation between cells in a cluster. We varied these two parameters independently. The former showed no significant change in the invasion modes, while the latter showed an increase in the amount of single cell migration for values between 0.001 and 0.075 but remained constant for higher values. For both parameters, the ratio between single cells and cells in clusters remains constant. Finally, the parameter *cell_ecm_repulsion* modulates the force of repulsion of the ECM that prevents the cells from moving across a voxel full of ECM and maintains them confined within the spheroid. The results of the analysis shows a significant drop in the amount of migrating single cells for values of repulsion higher than 20. For values between 5 and 20, the amount of single cells is constant and significantly higher than the number of cells in clusters. This is due to the fact that for low repulsion values, the cells can reach the ECM voxel more easily, and, with this contact, are able to change to a mesenchymal phenotype before developing cell-cell adhesion and junction.

## 3. Validation of the model by reproducing migration experiments

In order to validate the model, we successfully reproduced experimental observations reported in two studies: the first one focuses on the secretion of MMPs through the modification of p63 (41) and the second one on the local activation of SRC which leads to collective migration (33). These applications show how interventions to the intracellular or extracellular mechanisms of the cell can affect the behavior of the tumor growth and invasion. The model can then suggest means to interfere with these behaviors, as it is suggested for two of the examples. An additional example in the supplementary material considers the role of the ECM in the different modes of invasion (40).

### 3.1 Simulation of p63/MT1-MMP axis required for in situ to invasive transition

Lodillinsky and colleagues observed that, in basal-like breast cancer, the secretion of MT1-MMP and subsequent cell migration, was strictly linked to the up-regulation of p63 (41). It was shown in this study that the contact of cells with the ECM increased the level of an isoform of p63, Δp63 and as a consequence of MT1-MMP. Through the inhibition of p63, they noticed a strong decrease in the MT1-MMP level, thereby decreasing the process of invasion. The over-expression of MT1-MMP instead reestablished the invasive capacity of p63 depleted cells (Fig. 2, top panel). In this example, a modification of the intracellular model shows an impact at the population level. The node *MMPs* of the model accounts for a set of matrix metalloproteases consisting of MT1-MMP (MMP14), MMP13 and MMP2 (18) and its activity is regulated by Notch, SMAD, RAC1, p73 and p63 (see Supplementary Material for more details about the regulations). We first simulated the full inhibition of *p63* (setting the node to 0) in conditions that would activate the migration process (*Oxygen*=1, Growth_factor=1, Neighbor=1, ECM=1, TGF-beta=1, DNAdamage=0). In this simulation, the activation of MMPs was absent, blocking the invasive capacity of the cells and confining the tumor (Fig. 2 low panel). When p63 is over-expressed, and as a consequence MMPs are over-activated, the cells are able to degrade the ECM, allowing the tumor to expand and grow (Fig. 2) as it is observed in the initial experiment. The model was able to mimic experimental observations. We notice that the inhibition of p63 does not block the EMT activation. Here, the cells are still able to shift to a mesenchymal state, to grow and to divide. On the other hand, overexpression of p63 in the simulation promotes ECM degradation but blocks EMT transition, allowing tumor expansion but not single or collective migration as was reported by (41). To further explore this scenario, we searched for an additional mutation that could stop invasion in the p63/MMPs over-expression condition. We initially performed a sensitivity analysis of the intracellular model which consisted in automatically deleting and over-expressing all the nodes of the network (see Supplementary Materials), and selected ERK knock-out, which affects the expression of p21 and the phenotype *Cell_growth, Apoptosis* and *Migration*. We simulated the p63/MMPs conditions first, without the mutation, and we then introduced the knock-out halfway of the simulation. ERK knock-out caused the triggering of apoptosis in almost all the cells completely stopping the invasion process. Interestingly, the knock-out of ERK without p63 overexpression only caused a small percentage of cells to go to apoptosis without stopping the invasion process.

**Fig. 2.**
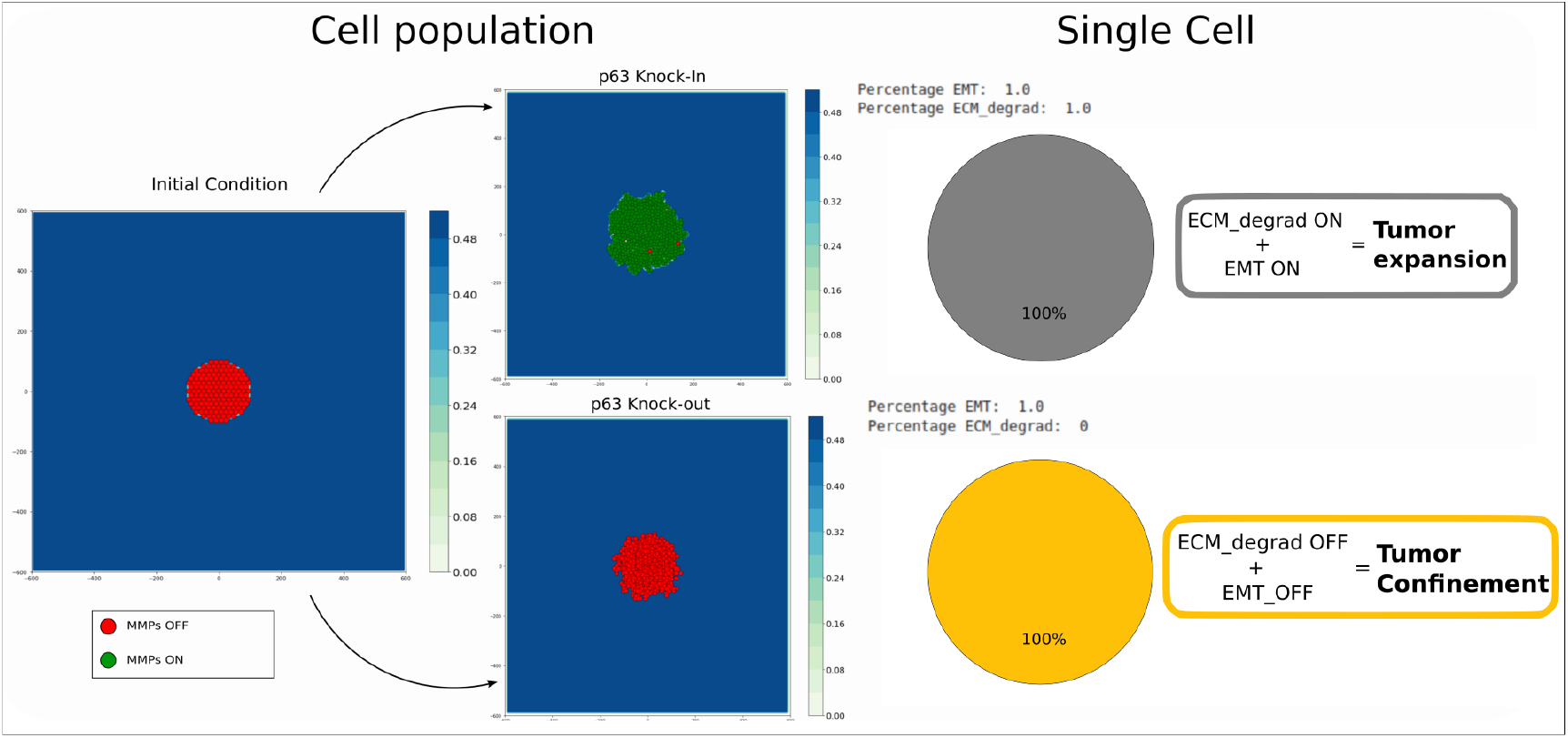
Reproduction of the p63 knock-in/knock-out experiment from (41). The initial conditions are the same for both simulations. In p63 overexpression condition, it is possible to verify a direct up-regulation of MMPs, leading to a ECM degradation phenotype. This is verified at the single cell level and is interpreted at the cell population level as tumor expansion. In p63 knock-out condition, it is possible to verify a down-regulation of MMPs, that lead to the inhibition of the ECM degradation phenotype. This is also verified at the single cell level and is interpreted at the cell population as tumor confinement.

### 3.2 Simulation of local light-activation of the SRC oncoprotein in an epithelial monolayer

Moitrier et al. used a synthetic light-sensitive version of SRC to trigger its activity upon light activation (33). They subjected a monolayer of cells to intermittent blue light to induce a spatially-constrained activation of the pro-invasive SRC protein and correlated this with the formation of extruded cells that remain cohesive. We adapted the model and used two different setups in 2 and 3D to simulate this experimental setting (see Methods). In the intracellular model, the SRC oncoprotein regulates the activity of CTNNB1 and CDH2 (42), CDH1 (30) and the production of vimentin (VIM) (43). SRC is activated by focal adhesion kinase (FAK). In the 2D scenario (Fig. 3), upon activation of the blue light, the affected cells start to develop a migratory phenotype similar to that of cells in contact with the ECM. SRC activation triggers the disgregation of cell junctions and EMT, pushing the cells to migrate towards the source of oxygen. The layer of epithelial cells surrounding the cells that have acquired the EMT phenotype confines them and favors the formation of aggregates that try to free themselves by pushing these EMT-like cells outwards, leading to collective migration. Removing the light source from monolayer reverse the cell’s phenotype. The cells that went through EMT return in the epithelial state, blocking the migration process. We then replicate the experiment in 3D (Fig. S4). For this case, the experiment setup was changed: the ECM was removed to avoid the activation of the mesenchymal phenotype at the border of the monolayer and placed the cells at the bottom of the domain. With this setup, the cells no longer push against the layer of epithelial cells, but rather migrate vertically forming an extrusion on top of the monolayer as observed in the original experiment. With the deactivation of the light source, the cells undergo a SRC inhibition, reversing the EMT phenotype and freezing. The epithelial cells in the simulation keep proliferating, filling the gap left by the mesenchymal cells. We extended the study by testing the effect of SRC overexpression on the whole tumor. As expected, it caused a burst of invasion, speeding up the formation of both clusters and migrating single cells. As in the previous example, we searched for a way to limit or even suppress the metastasis formation in these conditions. We first tried to stop the activation of the *ECM_degradation* phenotype through a knock-out of *p63*, which was insufficient to inhibit MMPs activation. After a parameter sensitivity analysis, we targeted the ERK gene. As in the previous example, we introduced an inhibitory mutation of ERK half-way through the simulation, which stopped the tumor progression not by promoting apoptosis, but rather by inhibiting the cell movement and cell cycle.

**Fig. 3.**
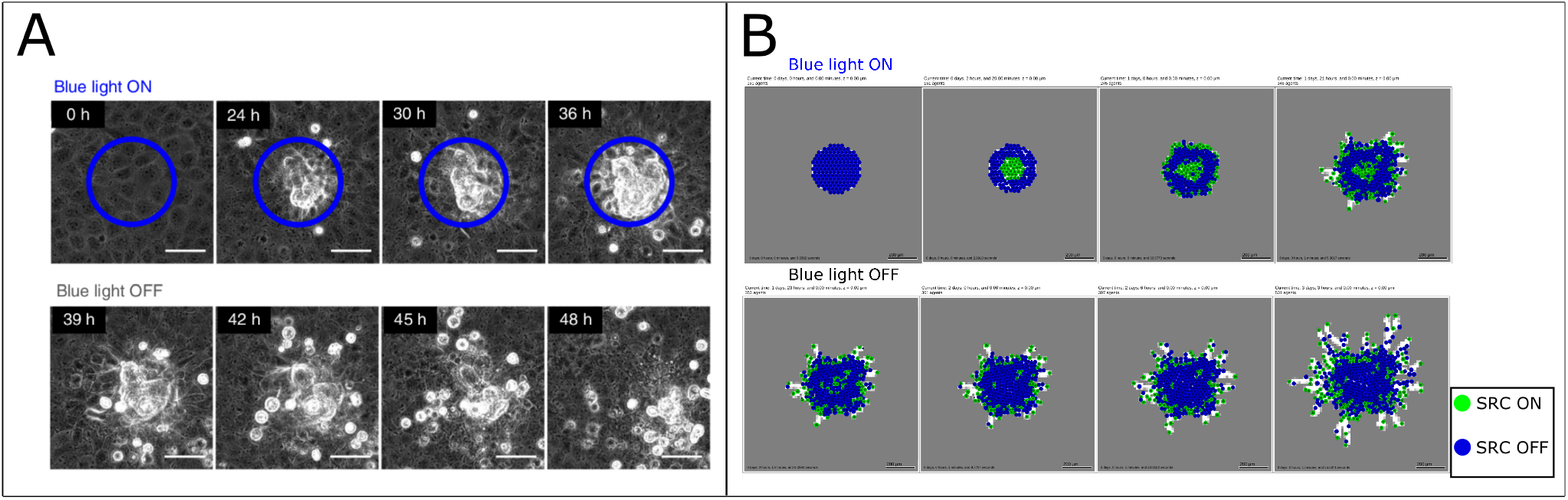
Reproduction of the SRC experiments. (A) Experimental results of SRC activation at different time points from (33). The blue circle indicates the area of the blue light activation. After 33 hours the light source is removed, reversing the collective extrusion phenotype. (B) Model simulation of the SRC experiments. We introduced a substrate that simulate the light in the middle of the epithelial monolayer (blue cells). The substrate virtually interacts with cells with a SRC activating mutation. The cells undergo EMT and become mesenchymal (green cells), trying to migrate and forming aggregates. When the substrate is removed, the mesenchymal cells return epithelials. SRC is found active at the borders of the monolayer in contact with ECM.

**Fig. 4.**
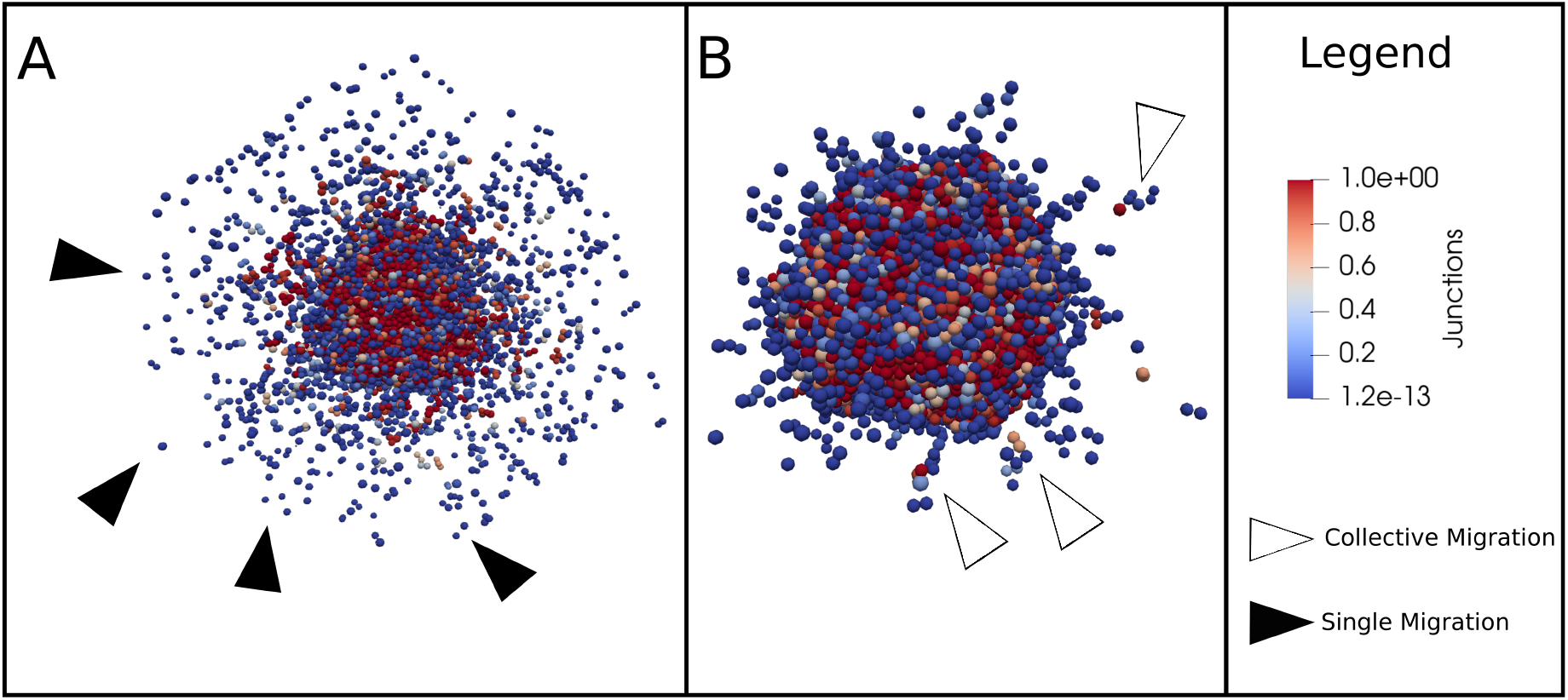
3D representation of the simulation. For panels A and B, different values of the parameters have been used to reproduce single (black arrows) and collective cell migration (white arrows). More details about the quantification of single and collective migration are given in the supplementary file, Section 5. The color bar represents the amount of cell junction used to establish cellular adhesion. (A) cell_ecm_repulsion = 10, epith_cell_junction_attach = 0.5, mes_cell_junction_detach = 0.03, migration_bias = 0.9, migration_speed = 0.8. (B) cell_ecm_repulsion = 15, cell_junction_attach = 0.005, cell_junction_detach = 0.001, migration_bias = 0.8, migration_speed = 0.7.

### 3.3 Simulation of drug candidates that can block migration

In addition to reproducing published studies, the model can also be used as a tool to explore means to revert invasive phenotypes or prioritize possible points of intervention and drug treatments with *in silico* experiments. We tested possible drug candidates that could block migration in conditions where cells have already undergone EMT and have started to invade the surrounding ECM. As a result of a systematic search for single and combined drug treatments, we found that overexpressing the activity of *CTNNB1* (forcing its value to 1) would not prevent tumor growth but would greatly limit the invasive capacity (Fig. S3). We tested this hypothesis by introducing an overexpression mutation both at the beginning and halfway through the simulation. In both cases, the treatment blocked the invasive capacities of the cells, preventing the activation of the node MMPs and, as a result, blocking the ECM degradation process. Identifying drugs that target CTNNB1 or any of its downstream actuators could have the potential of limiting invasion. In fact, there have been proposals for targeting CTNNB1 as the main player in the WNT canonical pathway in breast cancer as with 3,6-dihydroxyflavone in MDA-MB-231 cell line (44), even though many unknowns remain (45). A more extensive list of candidates can be found in the Jupyter notebook “intercellular_model_analysis”.

## 4. Discussion

In this study, we present a multiscale model that combines spatial cell representation and intracellular signaling to reproduce the different modes of cell invasion using PhysiBoSS. The model proved to be efficient in reproducing *in vitro* experiments and simulating different experimental scenarios. We managed to reproduce two invasion modes: single and collective, thanks to a combination of phenotypes and mechanical interactions with the ECM. This study confirms that tumor invasion is a complex process that benefits from considering spatial information, interaction with the microenvironment and intracellular representation

One application of this model could be to suggest and anticipate the potential risk of metastasis for patients that have a combination of mutations. Currently, the model includes a reasonable amount of genes to allow for a fast simulation and is able to capture the biological differences between the different modes of invasion. However, to be more precise in the model predictions, more genes and pathways could be included in the future based on new experiments. The choice of these pathways could be suggested by identifying differentially expressed genes or molecular signatures for patients with high metastatic potential, exploring public patient datasets such as The Cancer Genome Atlas (TCGA) data for which clinical data about tumor invasiveness are provided. In future releases, we also plan to include more cell types such as cancer associated fibroblast (CAFs) and T-cells to represent the role of the immune response. In this setting, each cell type will have its own intracellular model and would interact with other cell types according to biologically-relevant rules.

The current model has shown its predictive potential but there are several limitations. One of them is that the intracellular model can reproduce the EMT transition, but the commitment to EMT is final and non reversible. Nevertheless, we acknowledge the role of the mesenchymal to epithelial transition (MET) in the metastatis process which will investigate. The second limitation concerns the difficulty of choosing a proper set of parameters that can reproduce experimental data. While performing the sensitivity analysis, we mainly ran into two problems: 1) the number of parameters to analyze for this model is large and requires many long computations, and 2) we noted a high presence of noise in the results, even for a large number of simulations per parameter (50 runs for each tested value, see supplementary file, Section 5). This adds up to other computational costs of running PhysiBoSS simulations on a regular laptop. 2D simulations can take a couple of minutes, but 3D simulations require at least one hour. One way to tackle this issue is to run them on computer clusters with a high number of cores.

One way to address this parameter exploration issue is to rely on many individual simulations. For that, we plan to explore surrogate models and use them to learn subsets of parameters (46). More specifically, we want to train a machine learning algorithm connecting the inputs (parameter values) and the outputs of PhysiBoSS model, which will act as a surrogate for further parameter optimization. We will compare the output of the surrogate model with *in vitro*/*in vivo* data to find the best combination of parameters which will be tested in a relatively small number of simulations. With sufficiently many new simulations, the surrogate model will be synchronized with the original model by re-training to maintain the correspondence between the two models.

Finally, the multiscale model can be used as an exploratory and predictive tool to test hypotheses before performing wet lab experiments. The model is accompanied by the Jupyter Notebook and the Nanohub Tool to facilitate the reproducibility of the model results but also to allow users (biologists and/or modelers) to test additional experiments.

## ACKNOWLEDGEMENTS

We would like to thank Gaëlle Letort and Paul Macklin for inspiration and fruitful discussions.

This work has been partially supported by the European Commission under the PerMedCoE project (H2020-ICT-951773).

## 5. Supplementary Materials

Below is a list of supplementary file associated to this paper. It is possible to find the supplementary materials at https://github.com/sysbio-curie/Invasion_model_PhysiBoSS. It is possible to run the model online through the platform Nanohub: https://nanohub.org/resources/invasiongui.

- Supplementary File 1 - Supplementary information This document contains supplementary information and figures about the model, including a scheme to describe links of the variables of the agent-based model (ABM) model and of the Boolean model (BM), more information about the cell cycle in PhysiBoSS, a complete list of the logical formulae of the BM, a list of the main parameters of the ABM, the CTNNB1 overexpression simulation and the results of the sensitivity analysis.
- Supplementary File 2 - Intracellular_model_analysis In this jupyter notebook we provide a complete and detailed analysis of the intracellular model that we performed with MaBoSS.
- Supplementary File 3 - Variable_analysis In this jupyter notebook we provide a more detailed explanation on the evolution of the parameters that linked the intra-cellular model with the agent-based model.
- maboss_env.yml This file allows to build an environment using the conda tool, to run the jupyter notebook for the intracellulaer analysis.
- network.sif This file allows a fast representation of the intracellular model using Cytoscape
- intracellular_model.bnd This file contains the logical rules of the intracellular signalling network. It is essential in order to run both the model and the Supplementary File 2. It is in a format specific for MaBoSS.
- intracellular_model.cfg This file contains the initial configuration of the intracellular signalling network. It is essential in order to run both the model and the Supplementary File 2. It is in a format specific for MaBoSS.

