## Supplementary File 1 for "Multiscale model of the different modes of cancer cell invasion"

*********

### Multiscale modeling allows to study the different modes of cancer cell invasion

#### Outline

1. Connecting the agent-based model to the Boolean Model

2. Modeling with PhysiBoSS

3. EMT activation - molecular players

4. Cell Cycle model in PhysiBoSS

5. Experimental data used to validate the Boolean model

6. Simulation of initial conditions with MaBoSS

7. Logical formulae of the intracellular model

8. List of parameters of the model

9. CTNNB1 overexpressing mutation

10. Simulation of local light-activation of the SRC oncoprotein in an epithelial monolayer promotes collective extrusion - 3D scenario and ERK knock-out

11. Sensitivity analysis on model parameters

### 1. Connecting the agent-based model to the Boolean Model

We have described here the links of the variables of the agent-based model (ABM) model and of the Boolean model (BM).

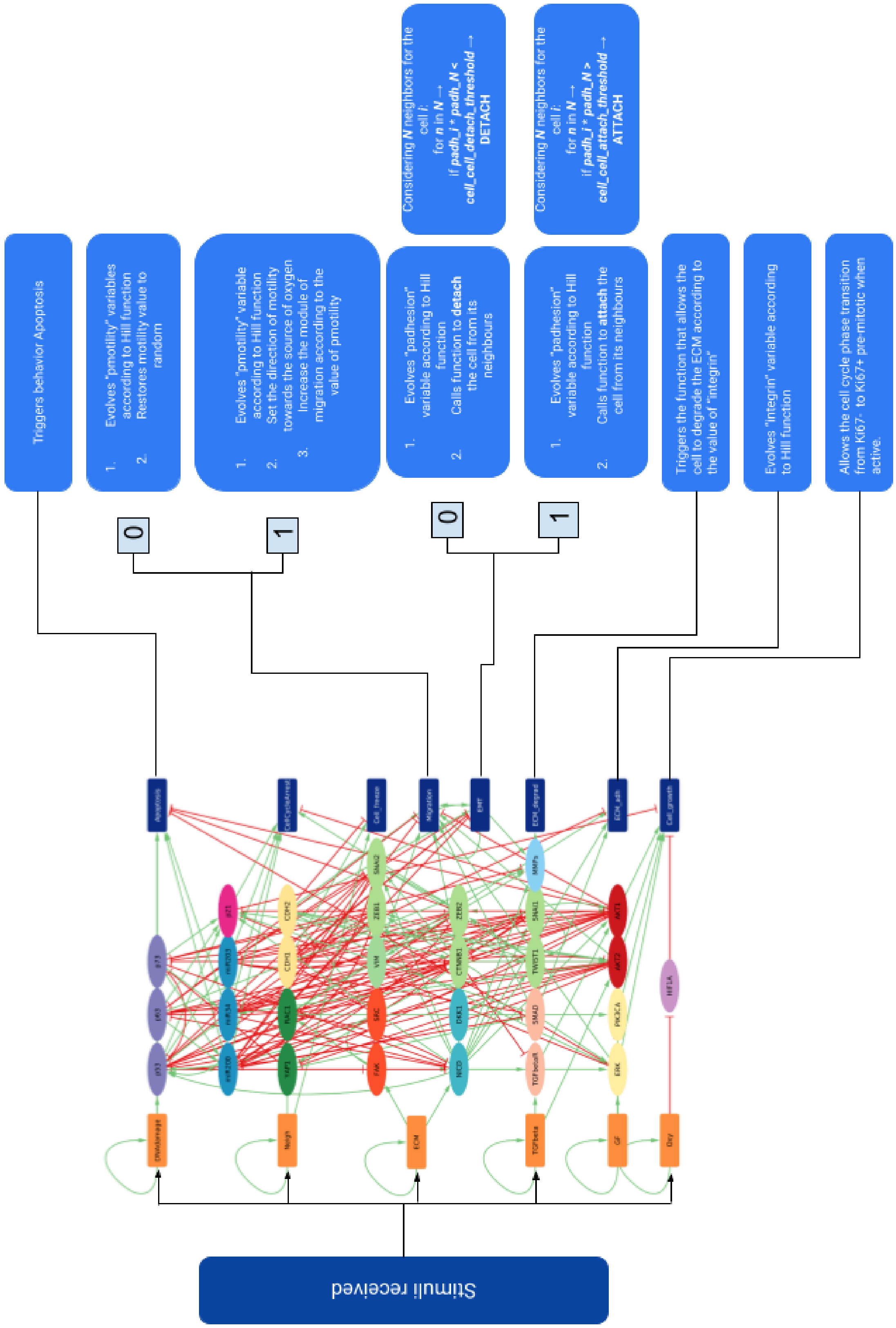

Figure S1: *Scheme of the links between the intracellular model and the behaviors of the agents.*

#### 2. Modeling with PhysiBoSS

Mutations or drug treatments are simulated by forcing the value of the corresponding nodes to 0 in the case of an inhibiting mutation (or treatment) and to 1 in the case of an activating mutation (or treatment).

We adapted the simulation set-ups to simulate the published experiments. For instance, to simulate the experiments from Moitrier and colleagues, we set up a one-cell thick circle of cells to obtain a monolayer of cells grown on a Petri dish. In addition, we introduced a parameter to represent the blue light activation. At a certain simulation time (measured in minutes), the blue light substrate appears in the simulation at the center of the monolayer, as it happens in the experiments.

#### 3. EMT activation - molecular players

EMT allows the cell to switch to a more motile phenotype, losing adhesion to neighboring cells and promoting invasiveness at the single cell level by reducing the expression of E-Cadherin (CDH1) and increasing the expression of N-Cadherin (CDH2). This cadherin switch is driven by the EMT regulators such as Twist1, Zeb1, Zeb2, Snai1, or Snai2 (Zhao et al. 2017) which activation can be induced by the focal adhesion kinase (FAK) activity, the tyrosine kinase SRC phosphorylation in response to cell-matrix contact (Hauck et al, 2002, Kerjouan et al. 2021), RhoGTPAses activity (RhoA, Rac1, Cdc42) (Parri et al 2010, Jung et al. 2020), or the presence of Transforming Growth Factor Beta (TGF-beta) (Plou et al. 2018).

#### 4. Cell Cycle model in PhysiBoSS

PhysiBoSS already provides the user with different options for cell cycle description.
For this study, we chose the model of the cell cycle based on Ki67 (Advanced Ki67 CellCycle model).

This cell cycle is composed of 3 phases and 3 transitions. Each cell starts in interphase Ki67-. Once the *Cell_growth* output node is activated, it triggers the transition to the next phase, Ki67+ pre-mitotic.

The cell then starts growing and once it reaches a certain threshold, it divides into two daughter cells that inherit the phenotype of the mother cell. Both daughter cells stay in Ki67+ post-mitotic phase and switch again to an early cell cycle phase Ki67- at a default transition rate provided by PhysiCell.

#### 5. Experimental data used to validate the Boolean model

We selected three publications with images and videos to validate the model. The first one is based on the work of Ilina et al. to explore how different densities of collagen, the major component of the ECM, affect the invasion modes. Figure S3 compares the simulated experiments with Figure 2.i of their work, where they used 4T1 breast cancer cell lines in 3D collagen matrices of different densities.

The second example compares the model to the experimental results of Lodillinsky and colleagues. In their work, they observed the regulation of MT1-MMP by upregulating p63 in breast cancer ductal carcinoma in situ (DCIS). The model is able to reproduce images presented in Figure of their publication where they explored the effects of the expression of a shRNA targeting p63 on invasion (immediately or after 2 days), as seen in Figure 2 in the main text. The configuration for this experiment involves multicellular spheroids of DCIS cells embedded in a 3D type I collagen matrix.

Finally, we considered the work of Moitrier et al. where it was shown that a direct activation of the SRC oncoprotein causes the displaying of some EMT activators in Madin-Darby Canine Kidney cells (MDCK). Their configuration involves a monolayer of MDCK cells that express a light-sensitive version of SRC. Using this approach, they were able to directly modulate the SRC expression using a blue light illumination pattern. We compare the results of our simulations with Moitrier's Figure 4.a, as seen in Figure 3 in the main text. The panel shows the cell flow and its reversibility during collective extrusion.

#### 6. Simulation of initial conditions with MaBoSS

The model of the individual cell can be simulated to study the different conditions surrounding the cell by setting the fixed inputs to values that represent the status of the microenvironment. Conditions at the center of the tumor or at the border of the tumor are not the same and the cell fates vary.

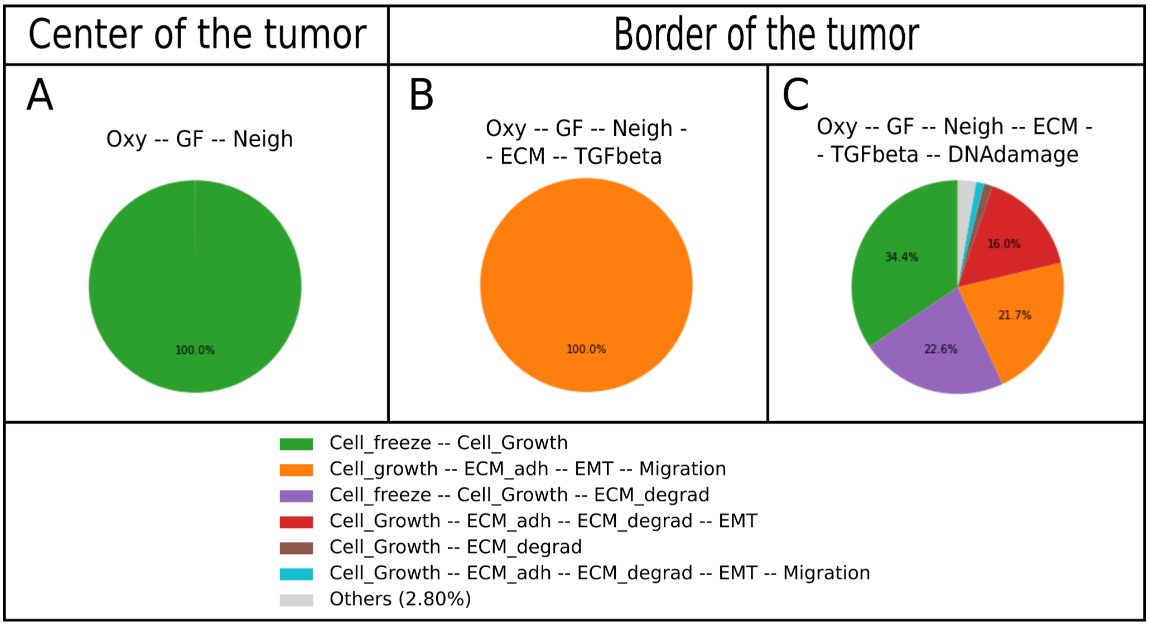

*Figure S2. Asymptotic solutions of the MaBoSS model are depicted as pie charts representing the proportion of cells in a state (i.e., phenotype). Two conditions are simulated: (A) at the center of the tumor with the presence of oxygen, growth factors and in contact with other cells, and with the same conditions but (B) at the border of the tumor, in contact with ECM and exposed to TGFbeta, and (C) with DNA damage.*

#### 7. Logical formulae of the intracellular model

The notation for the logical connectors is:
& for AND
| for OR
! for NOT

| Variable | Logical rule |
|---|---|
| HIF1A | !Oxy |
| FAK | (ECM \| (SRC)) & !p53 |
| YAP1 | ((!AKT1 \| ! AKT2) & SRC) |
| RAC1 | ( SRC \| FAK) & !(AKT1 \| AKT2 ) |
| PIK3CA | (GF \| RAC1) |
| MMPs | ( MMPs & (((NICD & SMAD)\| RAC1) & !p73) ) \| p63 |
| SRC | FAK |
| NICD | (!p73 & !p53 & !p63 & !miR34 & !miR200 & (ECM \| FAK)) |
| CTNNB1 | (!DKK1 & !p53 & !AKT1 & !p63 & !miR34 & !miR200 & !CDH1 & CDH2 & !SRC) |
| DKK1 | (!NICD & CTNNB1) \| (NICD) |
| AKT2 | TWIST1 & (TGFbeta \| GF \| CDH2) & !(miR203 \| miR34 \| p53) |
| ZEB1 | ((TWIST1 & SNAI1) \| CTNNB1 \| SNAI2 \| NICD) & ! miR200 |
| SNAI1 | (NICD \| TWIST1) & ! miR203 & ! miR34 & ! p53 & ! CTNNB1 |
| ZEB2 | (SNAI1 \| (SNAI2 & TWIST1) \| NICD) & ! miR200 & ! miR203 |
| p73 | (!AKT2 & !ZEB1 & !p53 & !AKT1 & DNAdamage & !YAP1) |
| p53 | (DNAdamage \| CTNNB1 \| NICD \| miR34) & ! SNAI2 & ! p73 & ! AKT1 & ! AKT2 |

| | |
|---|---|
| AKT1 | (CTNNB1 & (NICD \| TGFbetaR \| GF \| CDH2) & ! p53 & ! miR34 & ! CDH1) |
| p63 | (!NICD & !AKT2 & !p53 & !AKT1 & DNAdamage & !miR203) |
| miR34 | !(SNAI1 \| ZEB1 \| ZEB2) & (p53 \| p73) & AKT2 & ! p63 & ! AKT1 |
| SNAI2 | (TWIST1 \| CTNNB1 \| NICD) & ! miR200 & ! p53 & ! miR203 |
| miR200 | (p63 \| p53 \| p73) & !(AKT2 \| SNAI1 \| SNAI2 \| ZEB1 \| ZEB2) |
| TWIST1 | CTNNB1 \| NICD \| SNAI1 |
| CDH1 | (!AKT2 & !ZEB1 & !ZEB2 & !SNAI1 & !SNAI2 & !TWIST1 & !SRC & Neigh) |
| CDH2 | (TWIST1 \| SRC) |
| TGFbetaR | (NICD & !CTNNB1 & TGFbeta) |
| miR203 | (!ZEB1 & !ZEB2 & !SNAI1 & p53) |
| ERK | ((SMAD \| CDH2 \| GF \| NICD) & !AKT1) |
| SMAD | (!miR200 \| !miR203) & (TGFbetaR \| YAP1) |
| p21 | ((SMAD & NICD) \| p63 \| p53 \| p73 \| AKT2) & !(AKT1 \| ERK) |
| VIM | CTNNB1 \| ZEB2 \| SRC |
| EMT | (!CDH1 & CDH2) \| EMT & (!CDH1 & CDH2) |
| Migration | (AKT2 & !AKT1 & !miR200 & ERK & VIM & EMT & ((CDH2 & SMAD) \| (CTNNB1)) & !p63) |
| Apoptosis | (p53 \| p63 \| p73 \| miR200 \| miR34) & ! ZEB2 & ! AKT1 & ! ERK |
| ECM_adh | (NICD & !CDH1 & SMAD) \| RAC1 |
| ECM_degrad | MMPs |
| CellCycleArrest | (miR203 \| miR200 \| miR34 \| ZEB2 \| p21) & !AKT1 |
| Cell_freeze | (Neigh & !CDH2 & CDH1) |
| Cell_growth | ((ERK & !p21) \| (AKT1 & AKT2 & PIK3CA)) & !HIF1A |

#### 8. List of parameters of the model

Here we report a brief list of the main parameters of the simulation with a short description.

| Parameter | Description | Value |
|---|---|---|
| Domain | 3D / 2D space domain | 600x600(x600)μm |

| | | |
|---|---|---|
| ∆space | voxel’s unit measure | 10 µm |
| ***Cell-substrates interaction parameters*** | | |
| ecm_adhesion_min | set the min adhesion between cells and ECM | 1 |
| ecm_adhesion_max | set the max adhesion between cells and ECM | 2 |
| cell_ecm_repulsion | set the value of ECM repulsion | 15µm/min |
| ***Cell parameters*** | | |
| max_interaction_factor | set the max distance of interaction | 1.3µm |
| homotypic_adhesion_min | set the min adhesion between cells of the same type | 0.4 |
| homotypic_adhesion_max | set the max adhesion between cells of the same type | 0.8 |
| ***Threshold parameters*** | | |
| contact_ECM_threshold | change the threshold needed to trigger ECM interaction | 0.05 |
| contact_TGFβ_threshold | change the threshold needed to trigger TGFβ interaction | 0.02 |
| contact_cell_cell_threshold | change the threshold needed to trigger Neigh node | 0.3 |
| epith_cell_junctions_attach_threshold | change the threshold needed to attach cells in cluster with cell junction | 0.05 |
| mes_cell_junctions_detach_threshold | change the threshold needed to detach cells in cluster with cell junction | 0.03 |
| ***Motility parameters*** | | |
| migration_bias | change value of migration bias for cells with migration node active | 0.85 |
| migration_speed | change value of migration speed for cells with migration node active | 0.5µm/min |
| motility_amplitude_min | change the min value of motility amplitude | 0.1 |
| motility_amplitude_max | change the max value of motility amplitude | 0.8 |
| ***Substrates parameters*** | | |
| config_radius | change the initial radius of the tumor | 100µm |
| TGFβ_radius | change radius of the TGFβ substrate | 90µm |
| densityβ | change initial density of the TGFβ substrate | 0.4 |
| density_ECM | change initial density of the ECM substrate | 0.5 |
| ECM_degradation | change the amount of ECM degraded by the cells | 0.05 |
| ECM_TGFβ_ratio | change the amount of TGFβ degraded by the cells | 0.002 |
| TGFβ_degradation | change the threshold needed to start sensing TGFbeta inside a voxel with ECM | 0.75 |

#### 9. CTNNB1 overexpressing mutation

Following the analysis of the intracellular model (see Supplementary_file2-Intracellular_model_analysis) we tested a possible overexpressing mutation of CTNNB1. As shown in the figure, compared to the standard condition, this mutation is not preventing the tumor from growing, but greatly affects the invasive capacity.

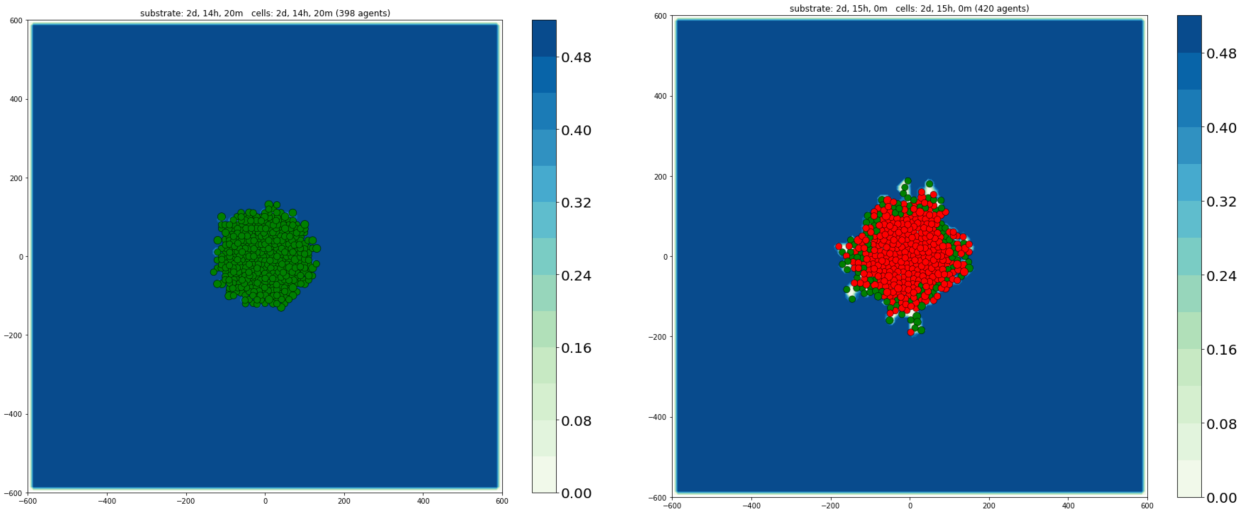

*Figure S3: On the left, simulation of the model in standard initial condition with no mutation. Green cells represent mesenchymal cells, red cells epithelial. On the right, knock-out mutation of CTNNB1.*

#### 10. Simulation of local light-activation of the SRC oncoprotein in an epithelial monolayer promotes collective extrusion - 3D scenario and ERK knock-out

The following are the 3D representation of the SRC experiment presented in 3.2 of the main text.

The different shades of red represent the amount of cell junctions. The cells in the center of the monolayer are affected by the light that causes the SRC activating mutation.

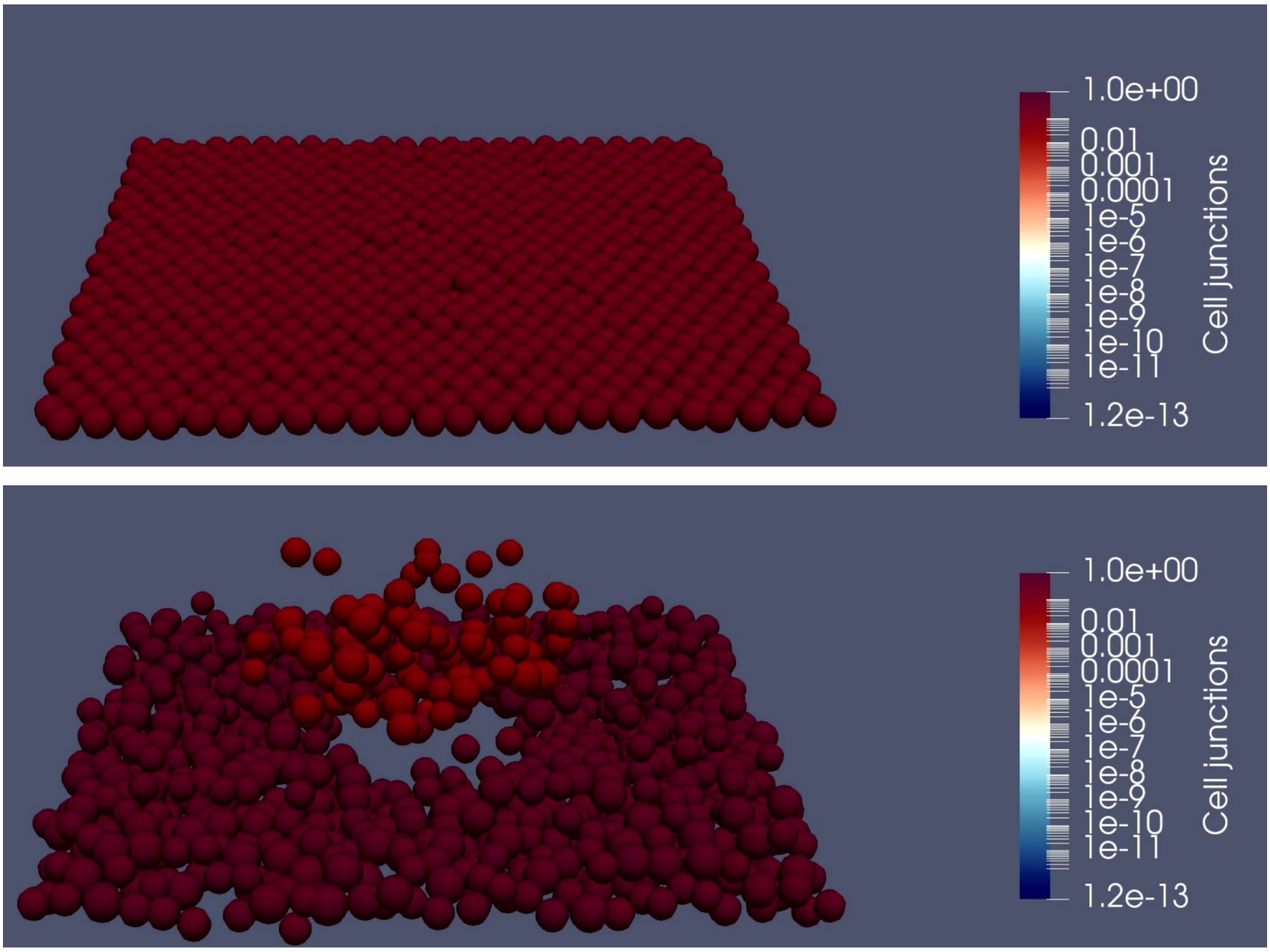

*Figure S4: 3D representation of the SRC in silico experiment. To avoid the activation of the mesenchymal phenotype, the extracellular matrix was removed. The cells at the center of the monolayer undergo EMT upon activation of SRC mutation.*

We then extended the 2D scenario simulating the SRC overexpression on the whole tumor. This caused a burst of invasion, speeding up the formation of both clusters and migrating single cells.
In the following snapshots, red cells are in an epithelial state, while green cells are in mesenchymal state.

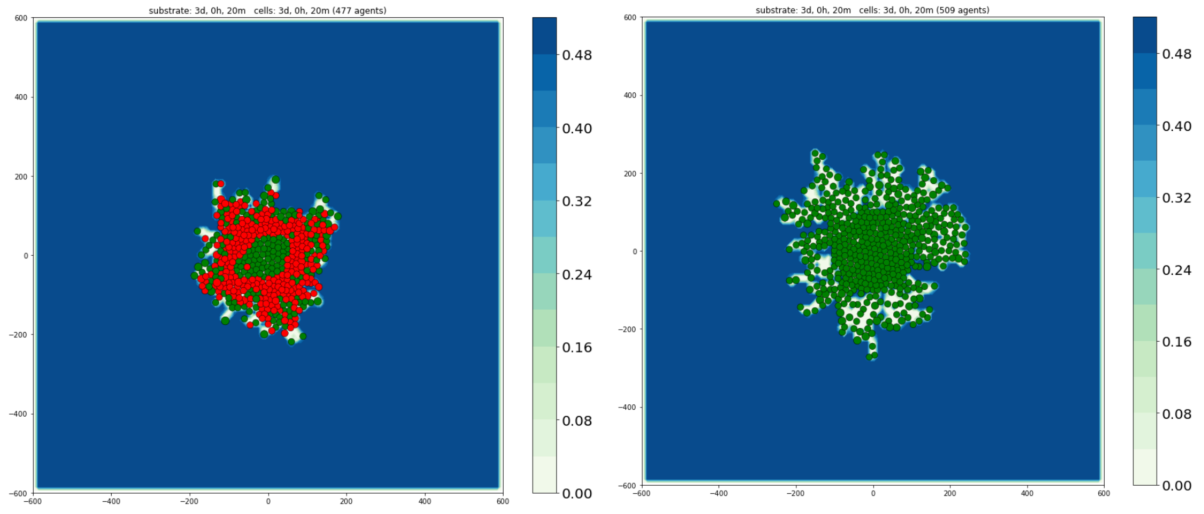

*Figure S5: comparison between SRC experiment with activating mutation in the middle of the monolayer (left panel) and at whole tumor level (right panel).*

As a further extension of the scenario, we simulated an ERK inhibition to try to stop the mesenchymal phenotype when SRC is overexpressed. We introduced mid simulation an ERK inhibiting mutation during the SRC overexpression on the whole tumor.

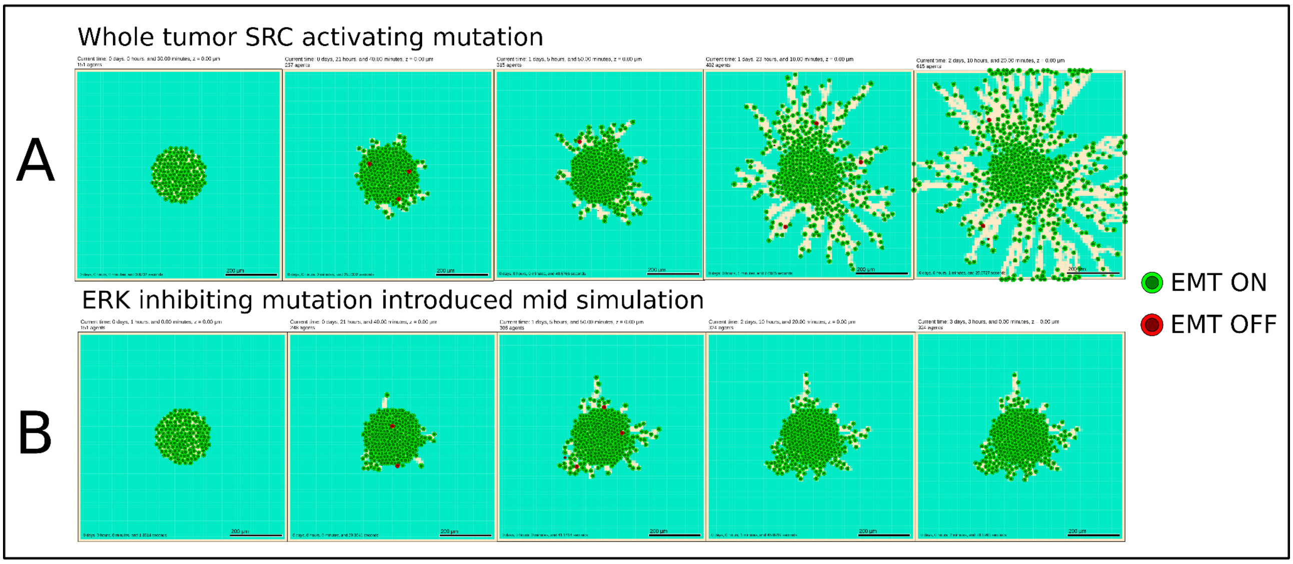

*Figure S6: A) simulation of SRC experiment with activating mutation at whole tumor level for different time points. B) Introduction of ERK knock-out mid simulation. In this case, ERK knock-out stop cell movement and proliferation without causing Apoptosis.*

#### 11. Cell adhesion and density changes in the ECM regulate modes of invasion

The first example explores the role of the ECM without any modification of the intracellular model. The model is able to reproduce images from data that were not used to build the model. In a recent study, Ilina and colleagues (Ilina et al. 2020) have shown how modifying the ECM density and uniformity affects the migration modes. The experiments were performed on the 4T1 breast cancer cell line, with partial expression of CDH1. The authors observed single cell detachment for low and medium density of the collagen, the experimental proxy of the ECM. In highly dense collagen, single cell invasion was no longer observed.
We tested the impact of changing the density of the ECM with our model by changing the value of the parameter *ECM_density.* In a uniform and highly dense ECM (*ECM_density* = 0.8), we confirm low single cell detachment and uncoordinated collective invasion (Figure S3-A). This is due to cells at the border of the tumor going through EMT, losing cell-junction adhesion, but remaining confined in the ECM. We further applied a uniform random distribution in the composition of the ECM of the voxels to simulate non-uniform low density ECM (0.1<*ECM_density* < 0.5). In this condition, we observed a high rate of single cell detachment as reported in the experiments (Figure S3-B). The model was able to reproduce both experimental observations and confirmed the role of the ECM in the types of migration that can occur during the early metastatic process.

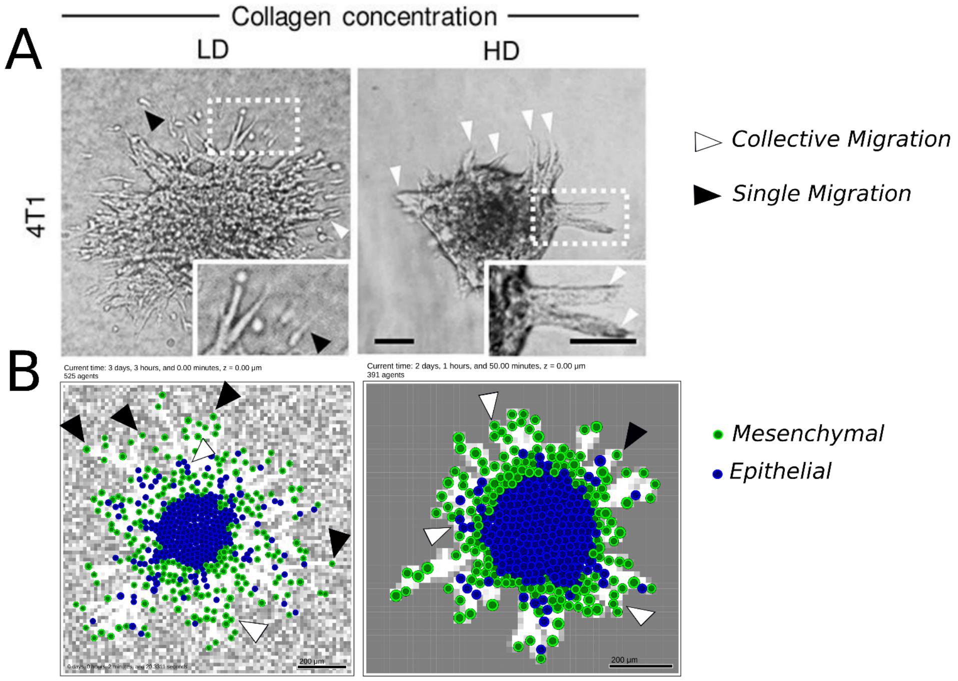

*Figure S7 [copyright from Ilina et al 2020]: Single vs. collective migration observations when varying ECM density and uniformity. (A) experimental observations in response to different ECM density (in uniform (right) and non-uniform conditions, reproduced from \citep{ilina_cellcell_2020}. (B) Model simulations of the uniform and high density ECM (right) and non-uniform low density ECM (left). White voxels represent low ECM density.*

#### 12. Sensitivity analysis on model parameters

We run a sensitivity analysis on the parameters of the ABM model that are linked to the BM. The goal is to measure how each parameter affects the amount of cells that migrate as single cells or as clusters. We select 7 parameters among the ones shown in the previous section. For each parameter, we choose a range of values to test based on our previous experience. For each value we did 50 runs and took the mean value and the squared mean error to see how the stochasticity of the simulation affected the results.

Each parameter has been tested independently. To limit the computational cost, we performed the sensitivity analysis on one parameter at a time.
The simulations took almost 90 hours on the cluster abacus at the Curie Institute (28 nodes for a total of 1120 cores, 5.25Tb of RAM).

To measure the amount of single and collective migration, we printed on a csv file the amount of interactive neighbors for each cell at each time step of the simulation. To separate the clusters and the single cells, we took advantage of a representation of the simulation as a network, using NetworkX to read the csv. Finally we counted the disconnected components of the resulting network, excluding the strongly connected component, which represents the core of the tumor.

The parameters are the following:

| **Parameters** | **Range** | **Potential range** | **Number of values selected** |
|---|---|---|---|
| *cell_ecm_repulsion*<br>regulates the amount of repulsion between cell and ECM | 5 < **15** < 50 | [0-infinite] | 10 |
| *epith_cell_attach_threshold*<br>changes the activation threshold needed to attach cells in cluster with cell junction | 0.001 < **0.05** < 1 | [0-1] | 25 |
| *mes_cell_detach_threshold*<br>change the activation threshold needed to detach cells in cluster with cell junction | 0.001 < **0.03** < 1 | [0-1] | 25 |
| *cell_cell_contact_threshold*<br>changes the activation threshold of the value *cell_contact* needed to trigger Neigh node | 0.01 < **0.3** < 3.5 | [0-infinite] | 18 |

| | | | |
|---|---|---|---|
| *cell_ecm_contact_threshold*<br>changes the activation threshold of the value *ecm_contact* needed to trigger ECM node | 0.001 < **0.05** < 1 | [0-1] | 27 |
| *migration_bias*<br>changes the value of migration bias for cells with *Migration* node active | 0.5 < **0.85** < 1 | [0-1] | 9 |
| *migration_speed*<br>changes the value of migration speed for cells with *Migration* node active | 0.3 < **0.7** < 1 | [0-1] | 7 |

The graphs below show the proportions of cells that are found as single cells (blue) or in clusters (orange) for various values of the parameters. Some results and interpretations are discussed in the main text.

- *epith_cell_attach_threshold*

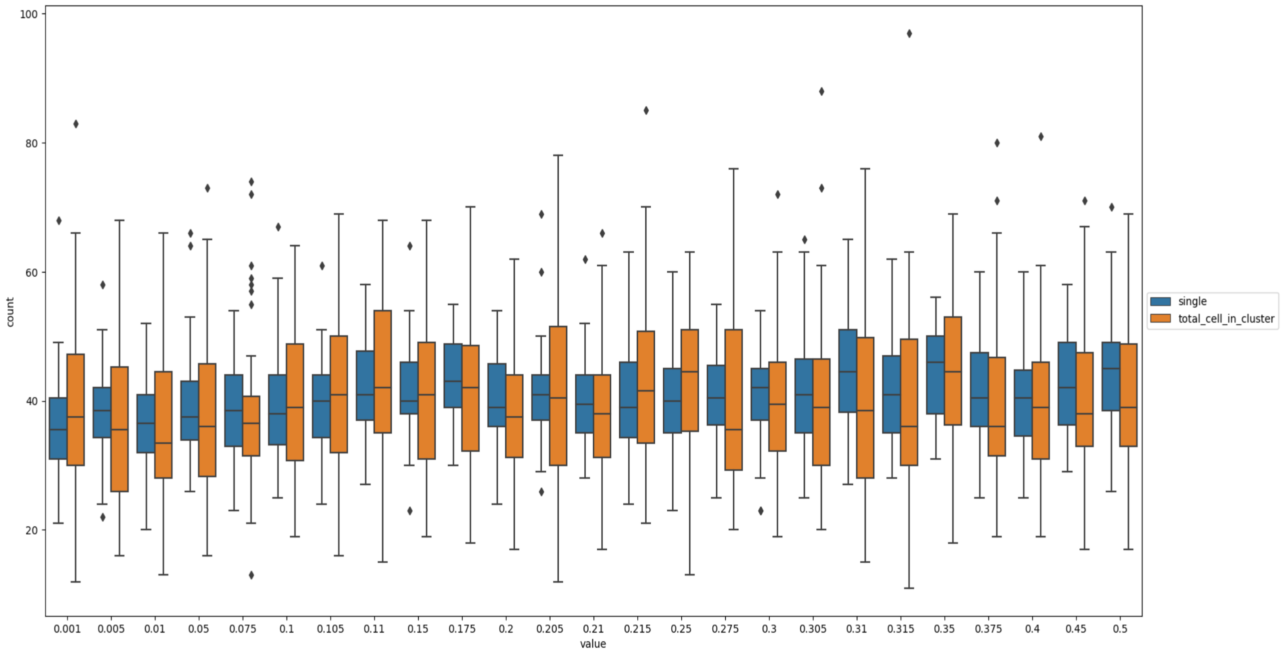

Figure S8: *Amount of single cell vs cell in cluster for different values of epith_cell_attach_threshold*

From the analysis, this parameter seems robust and shows a moderate change in the amount of single vs. collective migrating cells.

- *mes_cell_detach_threshold*

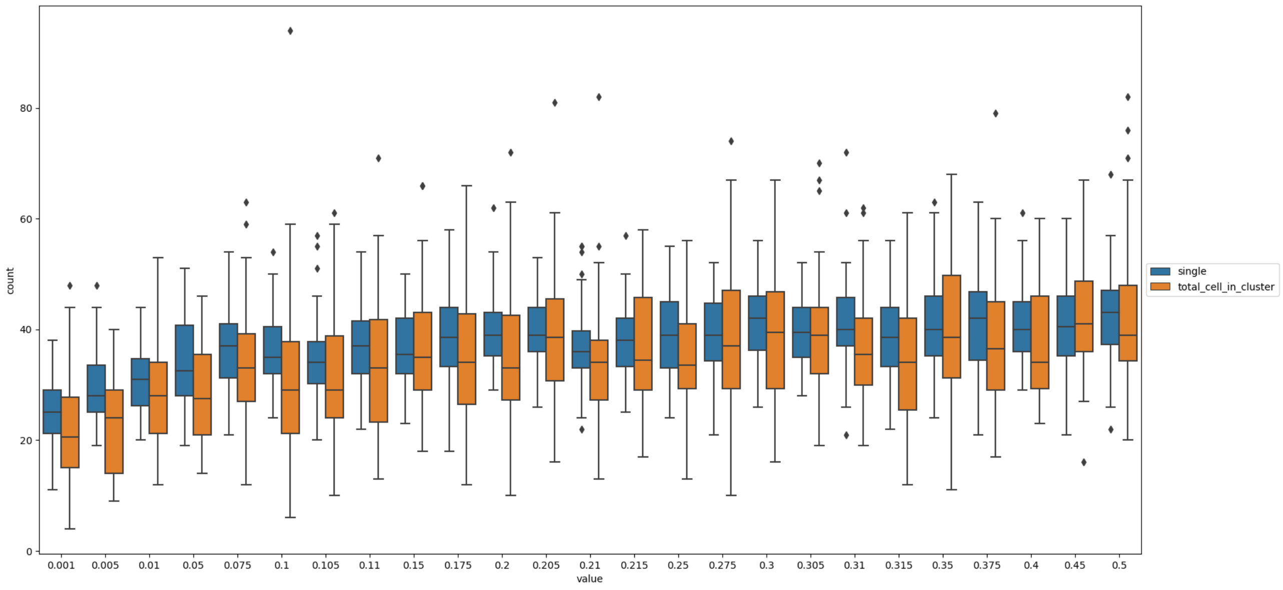

Figure S9: *Amount of single cell vs cell in cluster for different values of mes_cell_detach_threshold*

From the analysis, this parameter seems robust for values higher than 0.05 and it shows no changes in the rate between single and collective migrating cells. For values lower than 0.05, the amount of single vs. collective cells tends to diminish.

- *migration_bias*

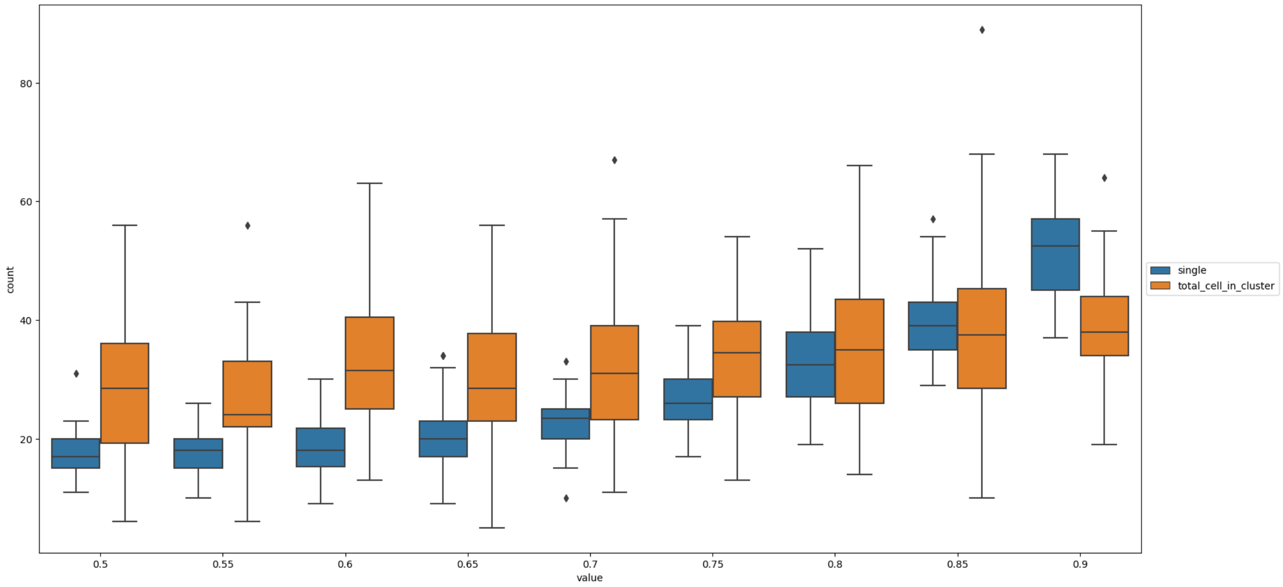

Figure S10: *Amount of single cell vs cell in cluster for different values of migration_bias*

From the analysis, this parameter heavily influences the amount of single and collective migration: for values minor than 0.85 the amount of cells migrating in clusters is higher than the single cells. For higher values, the number of single cells exponentially increases.

- *migration_speed*

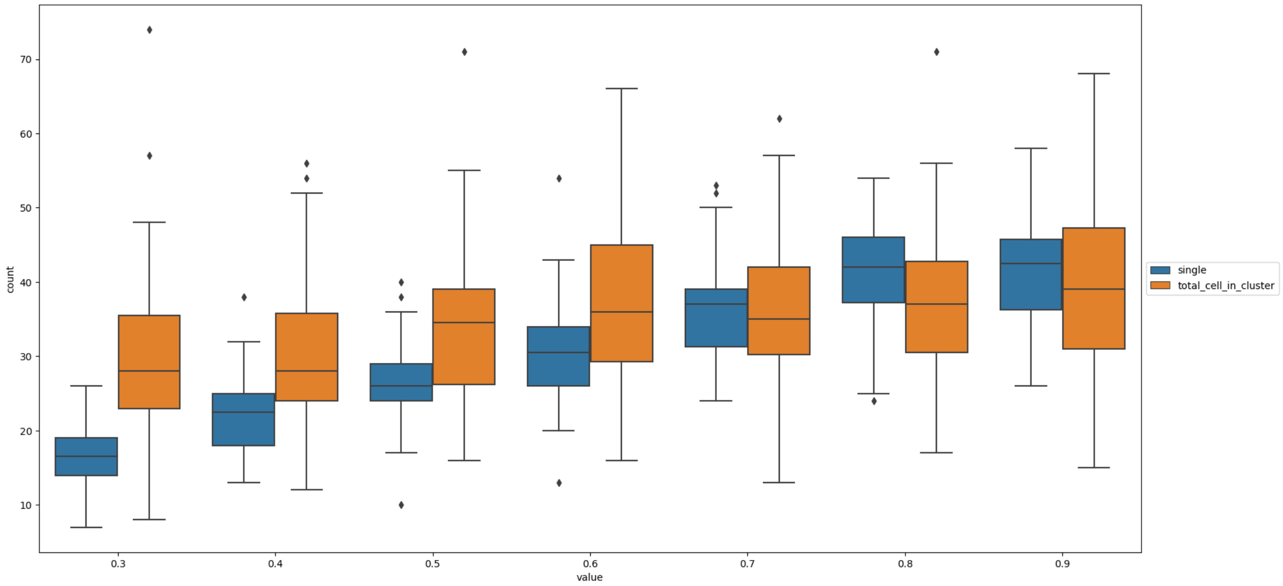

Figure S11: *Amount of single cell vs cell in cluster for different values of migration_speed*

From the analysis, this parameter heavily influences the amount of single and collective migration: for values minor than 0.7 the amount of cells migrating in clusters is higher than the single cells. For higher values, the number of single cells linearly increases.

- *cell_ecm_contact_threshold*

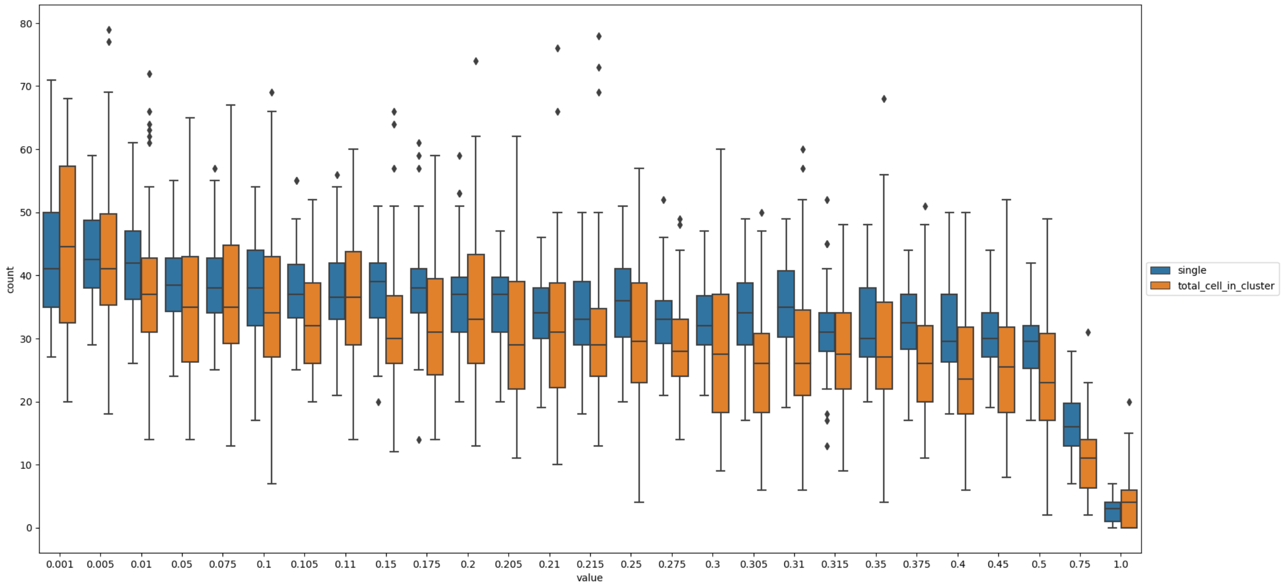

Figure S12: *Amount of single cell vs cell in cluster for different values of cell_ecm_contact_threshold*

From the analysis, this parameter seems to influence the amount of single and collective migration: the rate between single and collective migrating cells seems to be constant, decreasing slightly up to 0.5, after that it rapidly decreases.

- *cell_cell_contact_threshold*

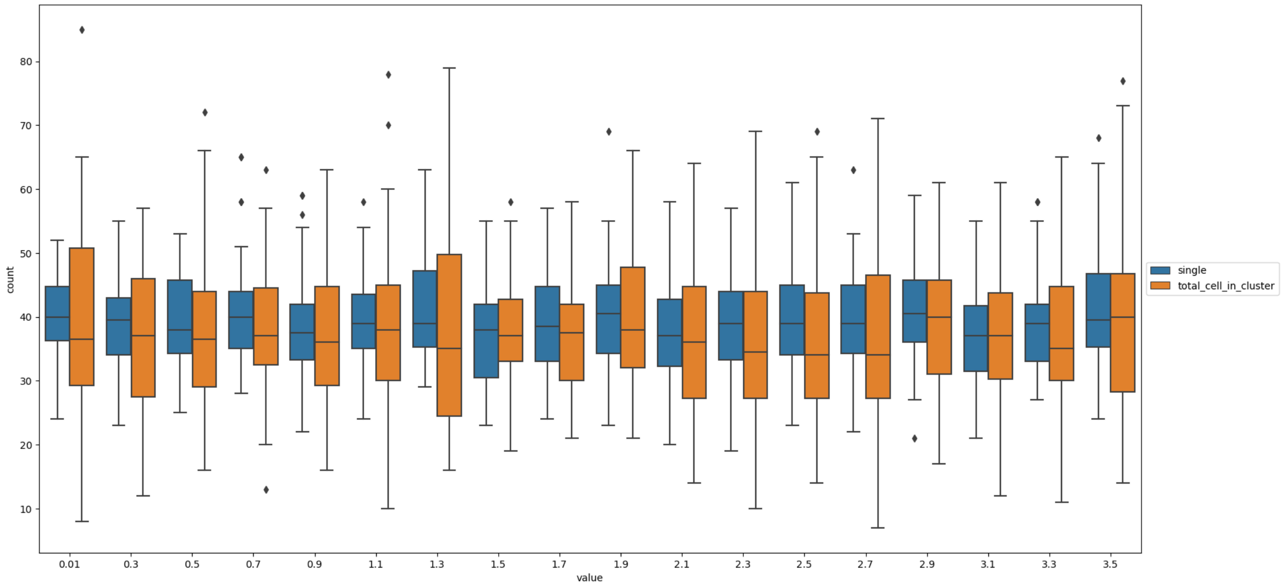

Figure S13: *Amount of single cell vs cell in cluster for different values of cell_cell_contact_threshold*

The analysis shows that the parameter has little impact on the separation of clusters and single cells.

- *cell_ecm_repulsion*

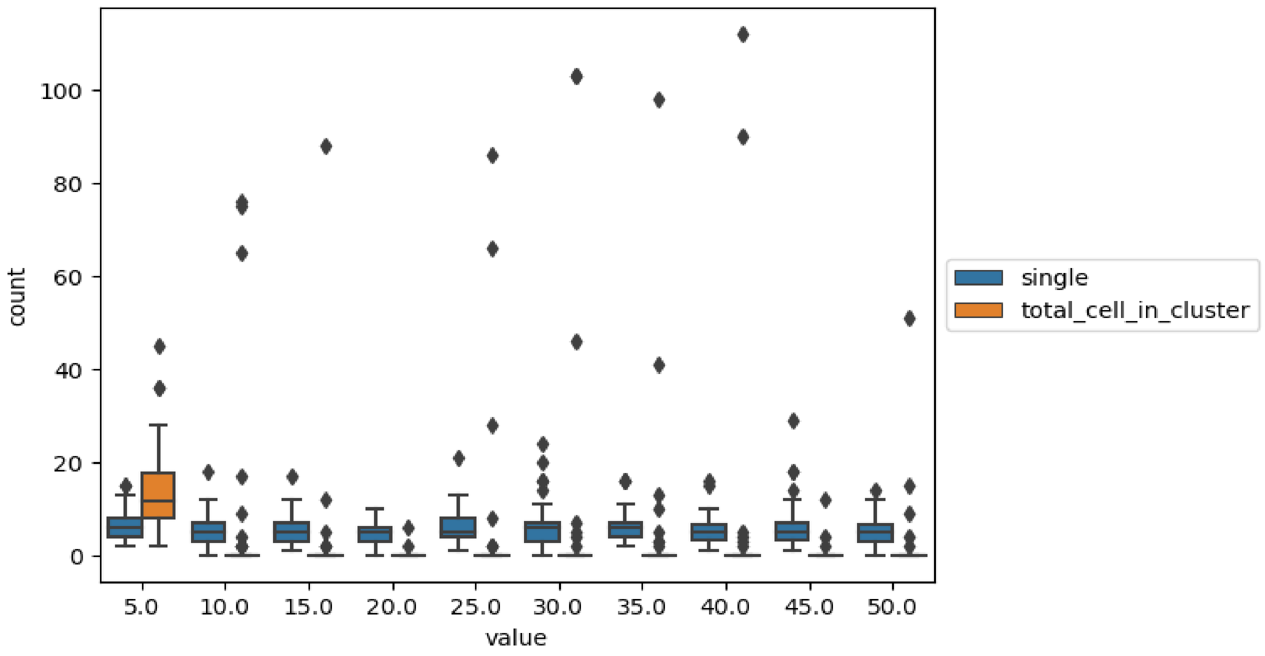

Figure S14: *Amount of single cell vs cell in cluster for different values of cell_ecm_repulsion*

With values higher than 5, it is difficult to see cells in clusters. This is due to the fact that, in these conditions, there are very few cells that touch the ECM and thus, that can become mesenchymal.
