## Supplementary File 2 for "Multiscale model of the different modes of cancer cell invasion"

Intracellular\_model\_analysis

This notebook is a companion for the article on a model describing the different modes of invasion using the tool PhysiBoSS.

PhysiBoSS is a modeling tool combining an agent-based approach for intercellular interactions and a Boolean model for the intracellular description.

We focus here on the Boolean model describing the signaling pathways inside each cell

#### Libraries for simulations¶

The libraries related to the tools needed for this analysis were developed in the context of CoLoMoTo, an initiative of the logical community to make all tools, methodologies and models in standard format to facilitate compatibility, interoperability and reproducibilty.

In [1]:

```
import maboss
import colomoto_jupyter
import maboss_test
import ginsim
import biolqm 
import boolean
import matplotlib.pyplot as plt
```

The Boolean model is simulated using MaBoSS, which is based on a stochastic simulation applied to the Boolean network. The MaBoSS model is composed of two files, one for the model description (.bnd) and the a second one for the definition of both the model and simulation parameters (.cfg).

In this notebook, we aim to show the behavior of the individual cell by fixing the values of the inputs that usually vary in the agent-based model. The purpose here is to explore the possible fates of the cell depending on the state of these inputs.

The analysis will first be done on the wild type case (with no alterations) in different initial conditions, which corresponds to the different conditions in which the cells may be found during the simulation. Next, we explore the cell dynamics in diseased conditions by simulating different gene mutations.

### Loading of the model¶

Here we load the .bnd and .cfg file necessary for the intracellular analysis.

The .bnd file contains the nodes and the logical formulas that regulates the interactions between genes. The .cfg file contains instead the parameters of the simulation for MaBoSS as well as the initial condition defined for the model.

We have defined a "standard initial condition", which represents a set of initial conditions for the model, defined in the MaBoSS configuration file ( .cfg provided in the supplementary materials).

The initial conditions are set as follows: ECM=0, TGFbeta=0, Neighbor=0, Oxygen=1, Growth\_factor=1, DNAdamage=0 (where 0 means inactive, 1 means active). Those conditions represent a favorable proliferating condition, where all the cells have access to oxygen and nutrients, are confined but not subjected to DNA damage or crowding.

Note that the simulation is set to represent the cells in an optimal condition to grow, and hence growth factors are assumed to be in a sufficient amount to constantly activate growth and maintain basal metabolism.

In [2]:

```
intracellular_model = maboss.load("./boolean_network/intracellular_model.bnd", "./boolean_network/intracellular_model.cfg")

on = [0, 1]
off = [1, 0]
```

### Network visualization¶

The network can be visualized using GINsim representation and exported into the standard format SBML-Qual.

In [11]:

```
def convert_and_show(model):   
    biolqm_model = maboss.to_biolqm(model)
    lrg = biolqm.to_ginsim(biolqm_model)
    return ginsim.show(lrg)
def convert_and_export(model):   
    biolqm_model = maboss.to_biolqm(model)
    biolqm.save(biolqm_model, "mymodel.bnet", "bnet")
    return
```

In [12]:

```
convert_and_export(intracellular_model)
convert_and_show(intracellular_model)
```

Out[12]:

### Definitions of the outputs of the model¶

The model outputs can be defined as a combination of some model node state.

In [4]:

```
set_output1 = ["Apoptosis", "Migration", "ECM_adh", "EMT", "Cell_growth", "ECM_degrad", "Cell_freeze"]
```

### Wild type simulation¶

To simulate the wild type conditions, we choose random initial values as there is no information about the initial state of the model entities.

In [5]:

```
intracellular_model.param["thread_count"] = 10
intracellular_model.param["max_time"] = 30
maboss.set_nodes_istate(intracellular_model, intracellular_model.network.keys(), {0: 0.5, 1: 0.5})
maboss.set_output(intracellular_model, set_output1)
```

In [6]:

```
res1 = intracellular_model.run()
res1.plot_piechart()
```

In [9]:

```
res1.plot_node_trajectory()
for phenotype in set_output1:
    try:
        print(phenotype, ": ", res1.get_last_nodes_probtraj()[phenotype].values[0])
    except:
        print(phenotype, ": ", 0)
```

```
Apoptosis :  0.13853
Migration :  0.54026
ECM_adh :  0.550113
EMT :  0.563881
Cell_growth :  0.392057
ECM_degrad :  0.416823
Cell_freeze :  0.213
```

#### Combinations of inputs: ECM/TGFbeta¶

We explore here the possible states of the cells at the border of the spheroid, which get in contact with the ECM and TGFbeta.

In [8]:

```
i_condition = intracellular_model.copy()
i_condition.param["max_time"] = 30
maboss.set_nodes_istate(i_condition, i_condition.network.keys(), off)
i_condition.network.set_istate("Oxy",on)
i_condition.network.set_istate("GF",on)
i_condition.network.set_istate("Neigh",on)
i_condition.network.set_istate("ECM",on)
i_condition.network.set_istate("TGFbeta",on)
i_condition.network.set_istate("DNAdamage",off)
maboss.set_output(i_condition, set_output1)
```

In [6]:

```
res2 = i_condition.run()
res2.plot_piechart()
```

In [12]:

```
res2.plot_node_trajectory()
for phenotype in set_output1:
    try:
        print(phenotype, ": ", res2.get_last_nodes_probtraj()[phenotype].values[0])
    except:
        print(phenotype, ": ", 0)
```

```
Apoptosis :  0
Migration :  1.0
ECM_adh :  1.0
EMT :  1.0
Cell_growth :  1.0
ECM_degrad :  0
Cell_freeze :  0
```

We explore now the phenotypes in the case of contact with ECM but in absence of Growth Factor

In [12]:

```
i_condition1_1 = intracellular_model.copy()
i_condition1_1.param["max_time"] = 30
maboss.set_nodes_istate(i_condition1_1, i_condition1_1.network.keys(), off)
i_condition1_1.network.set_istate("Oxy",on)
i_condition1_1.network.set_istate("GF",off)
i_condition1_1.network.set_istate("Neigh",on)
i_condition1_1.network.set_istate("ECM",on)
i_condition1_1.network.set_istate("TGFbeta",on)
i_condition1_1.network.set_istate("DNAdamage",off)
maboss.set_output(i_condition1_1, set_output1)
```

In [13]:

```
res2_1 = i_condition1_1.run()
res2_1.plot_piechart()
```

In [12]:

```
res2_1.plot_node_trajectory()
for phenotype in set_output1:
    try:
        print(phenotype, ": ", res2_1.get_last_nodes_probtraj()[phenotype].values[0])
    except:
        print(phenotype, ": ", 0)
```

```
Apoptosis :  0
Migration :  1.0
ECM_adh :  1.0
EMT :  1.0
Cell_growth :  1.0
ECM_degrad :  0
Cell_freeze :  0
```

We explore now the phenotypes in the case of contact with ECM but in absence of Growth Factor and with DNA damage

In [20]:

```
i_condition1_2 = intracellular_model.copy()
i_condition1_2.param["max_time"] = 60
maboss.set_nodes_istate(i_condition1_2, i_condition1_2.network.keys(), off)
i_condition1_2.network.set_istate("Oxy",on)
i_condition1_2.network.set_istate("GF",off)
i_condition1_2.network.set_istate("Neigh",on)
i_condition1_2.network.set_istate("ECM",on)
i_condition1_2.network.set_istate("TGFbeta",on)
i_condition1_2.network.set_istate("DNAdamage",on)
maboss.set_output(i_condition1_2, set_output1)
```

In [21]:

```
res2_2 = i_condition1_2.run()
res2_2.plot_piechart()
```

```
/home/marco/anaconda3/envs/MaBoSS_env/lib/python3.9/site-packages/maboss/figures.py:83: MatplotlibDeprecationWarning: normalize=None does not normalize if the sum is less than 1 but this behavior is deprecated since 3.3 until two minor releases later. After the deprecation period the default value will be normalize=True. To prevent normalization pass normalize=False 
  ax.pie(plot_line, labels=plotting_labels, radius=1.2,
```

In [22]:

```
res2_2.plot_node_trajectory()
for phenotype in set_output1:
    try:
        print(phenotype, ": ", res2_2.get_last_nodes_probtraj()[phenotype].values[0])
    except:
        print(phenotype, ": ", 0)
```

```
Apoptosis :  0.7648
Migration :  0.23049999999999998
ECM_adh :  0.23409999999999997
EMT :  0.23396399999999998
Cell_growth :  0.2352
ECM_degrad :  0.037073
Cell_freeze :  0.7656
```

#### Combinations of inputs: Oxy/GF¶

We explore here the possible states of the cells inside the spheroid, without contact with the ECM, but in optimal conditions to grow

In [8]:

```
i_condition2 = intracellular_model.copy()
i_condition2.param["max_time"] = 12
maboss.set_nodes_istate(i_condition2, i_condition2.network.keys(), off)
i_condition2.network.set_istate("Oxy",on)
i_condition2.network.set_istate("GF",on)
i_condition2.network.set_istate("Neigh",on)
i_condition2.network.set_istate("ECM",off)
i_condition2.network.set_istate("TGFbeta",off)
i_condition2.network.set_istate("DNAdamage",off)
maboss.set_output(i_condition2, set_output1)
```

In [9]:

```
res3 = i_condition2.run()
res3.plot_piechart()
```

In [15]:

```
res3.plot_node_trajectory()
for phenotype in set_output1:
    try:
        print(phenotype, ": ", res3.get_last_nodes_probtraj()[phenotype].values[0])
    except:
        print(phenotype, ": ", 0)
```

```
Apoptosis :  0
Migration :  0
ECM_adh :  0
EMT :  0
Cell_growth :  0.9999
ECM_degrad :  0
Cell_freeze :  1.0
```

#### Combinations of inputs: DNAdamage¶

We explore here the possible states of the cells at the border of the spheroid, in high pressure conditions (crowding), which cause nucleus deformation and as consequence, the rupture of nuclear envelope. This is then interpretated as DNA damage in the model.

In [13]:

```
i_condition3 = intracellular_model.copy()
i_condition3.param["max_time"] = 22
maboss.set_nodes_istate(i_condition3, i_condition3.network.keys(), off)
i_condition3.network.set_istate("Oxy",on)
i_condition3.network.set_istate("GF",on)
i_condition3.network.set_istate("Neigh",on)
i_condition3.network.set_istate("ECM",on)
i_condition3.network.set_istate("TGFbeta",on)
i_condition3.network.set_istate("DNAdamage",on)
maboss.set_output(i_condition3, set_output1)
```

In [14]:

```
res4 = i_condition3.run()
res4.plot_piechart()
```

```
/home/marco/anaconda3/envs/MaBoSS_env/lib/python3.9/site-packages/maboss/figures.py:83: MatplotlibDeprecationWarning: normalize=None does not normalize if the sum is less than 1 but this behavior is deprecated since 3.3 until two minor releases later. After the deprecation period the default value will be normalize=True. To prevent normalization pass normalize=False 
  ax.pie(plot_line, labels=plotting_labels, radius=1.2,
```

In [15]:

```
res4.plot_node_trajectory()
for phenotype in set_output1:
    try:
        print(phenotype, ": ", res4.get_last_nodes_probtraj()[phenotype].values[0])
    except:
        print(phenotype, ": ", 0)
```

```
Apoptosis :  0
Migration :  0.22949999999999998
ECM_adh :  0.408018
EMT :  0.397626
Cell_growth :  0.9999990000000001
ECM_degrad :  0.435288
Cell_freeze :  0.5812970000000001
```

The rupture of nuclear envelope, cause the activation of p63 that as conquence triggers the secretion of MMPs. This allows the degradation of the ECM and diminuish the pressure on the nucleus. Without DNA damage, the cell can undergo EMT and invade.

In the following case, we explore the effect of DNAdamage on the model in absence of growth factor (GF).

In [23]:

```
i_condition4 = intracellular_model.copy()
i_condition4.param["max_time"] = 12
maboss.set_nodes_istate(i_condition4, i_condition4.network.keys(), off)
i_condition4.network.set_istate("Oxy",off)
i_condition4.network.set_istate("GF",off)
i_condition4.network.set_istate("Neigh",on)
i_condition4.network.set_istate("ECM",off)
i_condition4.network.set_istate("TGFbeta",off)
i_condition4.network.set_istate("DNAdamage",on)
maboss.set_output(i_condition4, set_output1)
```

In [24]:

```
res5 = i_condition4.run()
res5.plot_piechart()
```

In [25]:

```
res5.plot_node_trajectory()
for phenotype in set_output1:
    try:
        print(phenotype, ": ", res5.get_last_nodes_probtraj()[phenotype].values[0])
    except:
        print(phenotype, ": ", 0)
```

```
Apoptosis :  1.0
Migration :  0
ECM_adh :  0
EMT :  0
Cell_growth :  0
ECM_degrad :  0.9211
Cell_freeze :  1.0
```

We notice how, in absence of GF, the model changed to an apoptotic phenotype. The growh factor input activate the ERK node in the intracellular model, which is necessary to substain the growth of the cells. In the model presented in the paper we decided to focus on the invasion aspect. For this reason, we run all the simulation in the best condition for the cells to grow.

### Mutants' simulation¶

#### CTNNB1 mutant¶

In [31]:

```
# Initial condition corresponds to i_condition2 defined in the wild type simulation
mutant2_simulation = maboss.copy_and_mutate(i_condition, ["CTNNB1"], "ON")

mut_res = mutant2_simulation.run()

mut_res.plot_piechart()

mut_res.plot_node_trajectory()
#mut_res.get_last_nodes_probtraj()["ECM_degrad"].values[0]
```

Conclusion: beta-catenin could be a potential target in the model to stop the release of MMPs and so the process of metastatisation. However, this mutation doesn't stop the EMT process and growth of the tumor.

#### TP63 mutants¶

##### TP63 overexpression¶

In [32]:

```
# Initial condition corresponds to a random initial condition as defined in the wild type simulation
mutant_simulation = maboss.copy_and_mutate(intracellular_model, ["p63"], "ON")
maboss.set_output(mutant_simulation, set_output1)

mut_res = mutant_simulation.run()

mut_res.plot_piechart()
```

In [33]:

```
mut_res.plot_node_trajectory()
for phenotype in set_output1:
    try:
        print(phenotype, ": ", mut_res.get_last_nodes_probtraj()[phenotype].values[0])
    except:
        print(phenotype, ": ", 0)
```

```
Apoptosis :  0.1056
Migration :  0
ECM_adh :  0.426972
EMT :  0.790937
Cell_growth :  0.45170000000000005
ECM_degrad :  1.0
Cell_freeze :  0.1048
```

Conclusion: overexpression of p63 on WT model increase the release of MMPs up to 100%, matching the results obtained from Lodillinski et al.

##### TP63 knock out¶

In [34]:

```
mutant_simulation = maboss.copy_and_mutate(intracellular_model, ["p63"], "OFF")
maboss.set_output(mutant_simulation, set_output1)

mut_res = mutant_simulation.run()

mut_res.plot_piechart()

emt = mut_res.get_last_nodes_probtraj()["EMT"].values[0]
degrad = mut_res.get_last_nodes_probtraj()["ECM_degrad"].values[0]

print("Percentage EMT: ", emt)
print("Percentage ECM_degrad: ", degrad)
```

```
Percentage EMT:  0.818936
Percentage ECM_degrad:  0.32920000000000005
```

```
/home/marco/anaconda3/envs/MaBoSS_env/lib/python3.9/site-packages/maboss/figures.py:83: MatplotlibDeprecationWarning: normalize=None does not normalize if the sum is less than 1 but this behavior is deprecated since 3.3 until two minor releases later. After the deprecation period the default value will be normalize=True. To prevent normalization pass normalize=False 
  ax.pie(plot_line, labels=plotting_labels, radius=1.2,
```

Conclusion: knock-out of p63 on WT model partially decrease the release of MMPs. To have a better understanding of the impact of overexpression of p63, we simulate next two different scenarios, specifying the initial condition for the model.

#### TP63 mutant with ECM/TGFbeta initial condition¶

##### TP63 overexpression¶

In [9]:

```
mutant_simulation = maboss.copy_and_mutate(i_condition, ["p63"], "ON")
maboss.set_output(mutant_simulation, set_output1)

mut_res = mutant_simulation.run()

mut_res.plot_piechart()

try:
    emt = mut_res.get_last_nodes_probtraj()["EMT"].values[0]
except:
    emt = 0

degrad = mut_res.get_last_nodes_probtraj()["ECM_degrad"].values[0]

print("Percentage EMT: ", emt)
print("Percentage ECM_degrad: ", degrad)
```

```
Percentage EMT:  1.0
Percentage ECM_degrad:  1.0
```

Conclusion: the initial condition applied for this case, simulate those cells at the border of the tumor in direct contact with the ECM. Overexpression of p63 leads totally leads to MMPs release which cause the degradation of the ECM. At the same time, p63 inhibits migration. This is in accord with the results from Lodillinski et al.

##### TP63 knock out¶

In [10]:

```
mutant_simulation = maboss.copy_and_mutate(i_condition, ["p63"], "OFF")
maboss.set_output(mutant_simulation, set_output1)

mut_res = mutant_simulation.run()

mut_res.plot_piechart()

try:
    emt = mut_res.get_last_nodes_probtraj()["EMT"].values[0]
except:
    emt = 0
try:
    degrad = mut_res.get_last_nodes_probtraj()["ECM_degrad"].values[0]
except:
    degrad = 0

print("Percentage EMT: ", emt)
print("Percentage ECM_degrad: ", degrad)
```

```
Percentage EMT:  1.0
Percentage ECM_degrad:  0
```

Conclusion: knock-out of p63 in this scenario, decrease the release of MMPs and allows the transition to mesenchymal state. This is partially in accord with the results from Lodillinsky. Knock-out of p63 should totally block MMPs release, confining the cells and avoiding metastatization.

#### SRC mutant¶

In this simulation, we test the effect of SRC activation on the model.

##### SRC overexpression¶

In [39]:

```
mutant_simulation5 = maboss.copy_and_mutate(intracellular_model, ["SRC"], "ON")

mut_res5 = mutant_simulation5.run()

mut_res5.plot_piechart()

print("Migration: ", mut_res5.get_last_nodes_probtraj()["Migration"].values[0])
print("EMT: ", mut_res5.get_last_nodes_probtraj()["EMT"].values[0])
```

```
Migration:  0.7667
EMT:  1.0
```

Conclusion: SRC overexpression in WT model increase the possibility for migration and EMT.

##### SRC knock out¶

In [40]:

```
mutant_simulation6 = maboss.copy_and_mutate(intracellular_model, ["SRC"], "OFF")

mut_res6 = mutant_simulation6.run()

mut_res6.plot_piechart()

print("Migration: ", mut_res6.get_last_nodes_probtraj()["Migration"].values[0])
print("EMT: ", mut_res6.get_last_nodes_probtraj()["EMT"].values[0])
```

```
Migration:  0.19160999999999997
EMT:  0.39782899999999993
```

Conclusion: Knock-out of SRC partially inhibits the formation of mesenchymal phenotypes. This is in accord with the fact that SRC is involved in many mechanisms, like adhesion, migration and degradation of ECM, forming complexes with FAK and promoting mesenchymal phenotype.

#### SRC mutant without ECM/TGFbeta initial condition (center of the monolayer)¶

##### SRC overexpression¶

In [41]:

```
mutant_simulation7 = maboss.copy_and_mutate(i_condition2, ["SRC"], "ON")

mut_res7 = mutant_simulation7.run()

mut_res7.plot_piechart()

print("Migration: ", mut_res7.get_last_nodes_probtraj()["Migration"].values[0])
print("EMT: ", mut_res7.get_last_nodes_probtraj()["EMT"].values[0])
```

```
Migration:  0.97828
EMT:  1.0
```

Conclusion:the results obtained are in accord with the experimental results shown in [ref]. Cells in the middle of the monolayer, upon activation of SRC, develop a mesenchymal-like phenotype.

##### SRC knock out¶

In [74]:

```
mutant_simulation8 = maboss.copy_and_mutate(i_condition2, ["SRC"], "OFF")

mut_res8 = mutant_simulation8.run()

mut_res8.plot_piechart()

#print(mut_res8.get_last_nodes_probtraj()["Migration"].values[0])
#print(mut_res8.get_last_nodes_probtraj()["EMT"].values[0])
```

Conclusion: knoc-out of SRC prevents the mesenchymal phenotype in cells in the middle of the layer.

### Model testing¶

In this section, we use a tool we developed, MaboSS\_test to verify that the model is able to reproduce observations or reported behaviors. These experimental observations are translated into a computer-readable format for fast-checking.

In [53]:

```
init_condition = {"Oxy":[0, 1], "Neigh":[0, 1], "GF":[0, 1], "DNAdamage":[1, 0], "ECM":[0, 1]}
```

In [49]:

```
p63_inhibition = {"p63":"OFF"}
p63_activation = {"p63":"ON"}
```

In [50]:

```
SRC_inhibition = {"SRC":"OFF"}
SRC_activation = {"SRC":"ON"}
```

In this simulation, we test that when p63 is overexpressed (or constitutively active), the probability of activating MMPs increase when compared to the wild type case.

For all the tests below the same format is applied:
[mutation to simulate, initial condition, observable (which read-out), behavior (increase, decrease)]

In [51]:

```
test_model = i_condition.copy()
run_test_model = maboss_test.MaBoSSTestCase(test_model)

run_test_model.assertStateProbabilityEvolution(p63_activation, init_condition, {"MMPs":1}, "increase")
```

```
True!  
The new probability of reaching the state is: 1.0 
The old one is: 0.248037
```

We then verify that the probability to activate MMPs decreases in the case of TP63 knock out.

In [52]:

```
test_model = i_condition.copy()
run_test_model = maboss_test.MaBoSSTestCase(test_model)

run_test_model.assertStateProbabilityEvolution(p63_inhibition, init_condition, {"MMPs":1}, "decrease")
```

```
True!  
The new probability of reaching the state is: 0 
The old one is: 0.241434
```

We then verify that the probability to activate EMT increase in the case of SRC overexpression.

In [55]:

```
test_model = i_condition2.copy()
run_test_model = maboss_test.MaBoSSTestCase(test_model)

run_test_model.assertStateProbabilityEvolution(SRC_activation, init_condition, {"EMT":1}, "increase")
```

```
True!  
The new probability of reaching the state is: 0.9999 
The old one is: 0.99604
```

We then verify that the probability to activate EMT decrease in the case of SRC knock-out.

In [57]:

```
test_model = i_condition2.copy()
run_test_model = maboss_test.MaBoSSTestCase(test_model)

run_test_model.assertStateProbabilityEvolution(SRC_inhibition, init_condition, {"EMT":1}, "decrease")
```

```
True!  
The new probability of reaching the state is: 0.990864 
The old one is: 0.995688
```

We then verify that the probability to activate Migration increase in the case of SRC overexpression.

In [59]:

```
test_model = i_condition2.copy()
run_test_model = maboss_test.MaBoSSTestCase(test_model)

run_test_model.assertStateProbabilityEvolution(SRC_activation, init_condition, {"Migration":1}, "increase")
```

```
True!  
The new probability of reaching the state is: 0.989715 
The old one is: 0.970647
```

We then verify that the probability to activate Migration decrease in the case of SRC knock-out.

In [60]:

```
test_model = i_condition2.copy()
run_test_model = maboss_test.MaBoSSTestCase(test_model)

run_test_model.assertStateProbabilityEvolution(SRC_inhibition, init_condition, {"Migration":1}, "decrease")
```

```
True!  
The new probability of reaching the state is: 1.6e-05 
The old one is: 0.972376
```

Conclusion: we verified that in the intracellular model, p63 directly control the activation of MMPs, proving to be a good candidate to stop tumor progression, inhibiting the release of matrix metalloproteases.
On the other hand, overexpression of SRC leads those cells in the middle of the monolayer, in an epithelial state, to differentiate into mesenchymal phenotype. Knock-out of SRC instead, block this differentiation, preventing the switch to mesenchymal phenotype.

### Automatic (single/double) mutations¶

Here we perform a sensititvity analysis on the model, triggering one or two mutations at the time over the nodes on the network. The purpose is to search for single or double mutants that exhibit interesting phenotypes, such as complete loss of migration, apoptosis, etc.

In [61]:

```
def simulate_single_mutants(model, list_nodes, sign="BOTH"):
    
    list_single_mutants = []
    if (sign == "BOTH" or sign == "ON"):
        list_single_mutants += [(node, "ON") for node in list_nodes]
    if (sign == "BOTH" or sign == "OFF"):
        list_single_mutants += [(node, "OFF") for node in list_nodes]
    
    res = {}
    for single_mutant in list_single_mutants:
        t_model = model.copy()
        t_model.mutate(*single_mutant)
        res.update({single_mutant: t_model.run()})
        
    return res
```

In [62]:

```
def simulate_double_mutants(model, list_nodes, sign="BOTH"):
    
    list_single_mutants = []
    if (sign == "BOTH" or sign == "ON"):
        list_single_mutants += [(node, "ON") for node in list_nodes]
    if (sign == "BOTH" or sign == "OFF"):
        list_single_mutants += [(node, "OFF") for node in list_nodes]
    
    list_double_mutants = [(a, b) for idx, a in enumerate(list_single_mutants) for b in list_single_mutants[idx + 1:] if a[0] != b[0]]

    res = {}
    for double_mutant in list_double_mutants:
        t_model = model.copy()
        t_model.mutate(*(double_mutant[0]))
        t_model.mutate(*(double_mutant[1]))
        res.update({double_mutant: t_model.run()})
        
    return res
```

In [63]:

```
def filter_sensititivy(results, state=None, node=None, minimum=None, maximum=None):
    ret_res = {}
    for (mutant, res) in results.items():
        
        if state is not None:
            t_res = res.get_last_states_probtraj()
            if state in t_res.columns:
                if minimum is not None and maximum is not None:
                    if t_res[state].values[0] > minimum and t_res[state].values[0] < maximum:
                        ret_res.update({mutant: res})
                elif minimum is not None and t_res[state].values[0] > minimum:
                    ret_res.update({mutant: res})
                elif maximum is not None and t_res[state].values[0] < maximum:
                    ret_res.update({mutant: res})
            elif maximum is not None and minimum is None:
                ret_res.update({mutant:res})

        elif node is not None:
            
            t_res = res.get_last_nodes_probtraj()

            if node in t_res.columns:
                if minimum is not None and maximum is not None:
                    if t_res[node].values[0] > minimum and t_res[node].values[0] < maximum:
                        ret_res.update({mutant: res})
                elif minimum is not None and t_res[node].values[0] > minimum:
                    ret_res.update({mutant: res})
                elif maximum is not None and t_res[node].values[0] < maximum:
                    ret_res.update({mutant: res})
            elif maximum is not None and minimum is None:
                ret_res.update({mutant: res})

    return ret_res
```

In [64]:

```
intracellular_model.network.keys()

target_genes = ["AKT1", "CTNNB1", 'DKK1', 'AKT2', 'RAC1', 'YAP1', 'PIK3CA', 'ERK', 'p21', 'SMAD', 'VIM', 'p53'
               , 'p63', 'p73', 'miR203', 'miR34', 'miR200', 'TGFbetaR', 'FAK']
```

In [65]:

```
i_condition4 = intracellular_model.copy()
i_condition4.param["max_time"] = 30
maboss.set_nodes_istate(i_condition4, i_condition4.network.keys(), off)
i_condition4.network.set_istate("Oxy",on)
i_condition4.network.set_istate("GF",on)
i_condition4.network.set_istate("Neigh",off)
i_condition4.network.set_istate("ECM",on)
i_condition4.network.set_istate("TGFbeta",on)
i_condition4.network.set_istate("DNAdamage",off)
maboss.set_output(i_condition4, set_output1)
```

In [66]:

```
res_single = simulate_single_mutants(i_condition, target_genes)
for mutant, res in res_single.items():
    print(mutant)
    res.plot_piechart()
    plt.title(mutant)
```

```
('AKT1', 'ON')
('CTNNB1', 'ON')
('DKK1', 'ON')
('AKT2', 'ON')
('RAC1', 'ON')
('YAP1', 'ON')
('PIK3CA', 'ON')
('ERK', 'ON')
('p21', 'ON')
('SMAD', 'ON')
('VIM', 'ON')
('p53', 'ON')
('p63', 'ON')
('p73', 'ON')
('miR203', 'ON')
('miR34', 'ON')
('miR200', 'ON')
('TGFbetaR', 'ON')
('FAK', 'ON')
('AKT1', 'OFF')
('CTNNB1', 'OFF')
('DKK1', 'OFF')
('AKT2', 'OFF')
('RAC1', 'OFF')
('YAP1', 'OFF')
('PIK3CA', 'OFF')
('ERK', 'OFF')
('p21', 'OFF')
('SMAD', 'OFF')
('VIM', 'OFF')
('p53', 'OFF')
('p63', 'OFF')
('p73', 'OFF')
('miR203', 'OFF')
('miR34', 'OFF')
('miR200', 'OFF')
('TGFbetaR', 'OFF')
('FAK', 'OFF')
```

```
/home/marco/anaconda3/envs/MaBoSS_env/lib/python3.9/site-packages/maboss/figures.py:83: MatplotlibDeprecationWarning: normalize=None does not normalize if the sum is less than 1 but this behavior is deprecated since 3.3 until two minor releases later. After the deprecation period the default value will be normalize=True. To prevent normalization pass normalize=False 
  ax.pie(plot_line, labels=plotting_labels, radius=1.2,
/home/marco/anaconda3/envs/MaBoSS_env/lib/python3.9/site-packages/maboss/results/baseresult.py:100: RuntimeWarning: More than 20 figures have been opened. Figures created through the pyplot interface (`matplotlib.pyplot.figure`) are retained until explicitly closed and may consume too much memory. (To control this warning, see the rcParam `figure.max_open_warning`).
  self._piefig, self._pieax = plt.subplots(1, 1)
```

In [67]:

```
res_double = simulate_double_mutants(i_condition, target_genes)
for mutant, res in res_double.items():
    print(mutant)
    res.plot_piechart()
    plt.title(mutant)
```

```
(('AKT1', 'ON'), ('CTNNB1', 'ON'))
(('AKT1', 'ON'), ('DKK1', 'ON'))
(('AKT1', 'ON'), ('AKT2', 'ON'))
(('AKT1', 'ON'), ('RAC1', 'ON'))
(('AKT1', 'ON'), ('YAP1', 'ON'))
(('AKT1', 'ON'), ('PIK3CA', 'ON'))
(('AKT1', 'ON'), ('ERK', 'ON'))
(('AKT1', 'ON'), ('p21', 'ON'))
(('AKT1', 'ON'), ('SMAD', 'ON'))
(('AKT1', 'ON'), ('VIM', 'ON'))
(('AKT1', 'ON'), ('p53', 'ON'))
(('AKT1', 'ON'), ('p63', 'ON'))
(('AKT1', 'ON'), ('p73', 'ON'))
(('AKT1', 'ON'), ('miR203', 'ON'))
(('AKT1', 'ON'), ('miR34', 'ON'))
(('AKT1', 'ON'), ('miR200', 'ON'))
(('AKT1', 'ON'), ('TGFbetaR', 'ON'))
(('AKT1', 'ON'), ('FAK', 'ON'))
(('AKT1', 'ON'), ('CTNNB1', 'OFF'))
(('AKT1', 'ON'), ('DKK1', 'OFF'))
(('AKT1', 'ON'), ('AKT2', 'OFF'))
(('AKT1', 'ON'), ('RAC1', 'OFF'))
(('AKT1', 'ON'), ('YAP1', 'OFF'))
(('AKT1', 'ON'), ('PIK3CA', 'OFF'))
(('AKT1', 'ON'), ('ERK', 'OFF'))
(('AKT1', 'ON'), ('p21', 'OFF'))
(('AKT1', 'ON'), ('SMAD', 'OFF'))
(('AKT1', 'ON'), ('VIM', 'OFF'))
(('AKT1', 'ON'), ('p53', 'OFF'))
(('AKT1', 'ON'), ('p63', 'OFF'))
(('AKT1', 'ON'), ('p73', 'OFF'))
(('AKT1', 'ON'), ('miR203', 'OFF'))
(('AKT1', 'ON'), ('miR34', 'OFF'))
(('AKT1', 'ON'), ('miR200', 'OFF'))
(('AKT1', 'ON'), ('TGFbetaR', 'OFF'))
(('AKT1', 'ON'), ('FAK', 'OFF'))
(('CTNNB1', 'ON'), ('DKK1', 'ON'))
```

```
/home/marco/anaconda3/envs/MaBoSS_env/lib/python3.9/site-packages/maboss/results/baseresult.py:100: RuntimeWarning: More than 20 figures have been opened. Figures created through the pyplot interface (`matplotlib.pyplot.figure`) are retained until explicitly closed and may consume too much memory. (To control this warning, see the rcParam `figure.max_open_warning`).
  self._piefig, self._pieax = plt.subplots(1, 1)
```

```
(('CTNNB1', 'ON'), ('AKT2', 'ON'))
(('CTNNB1', 'ON'), ('RAC1', 'ON'))
(('CTNNB1', 'ON'), ('YAP1', 'ON'))
(('CTNNB1', 'ON'), ('PIK3CA', 'ON'))
(('CTNNB1', 'ON'), ('ERK', 'ON'))
(('CTNNB1', 'ON'), ('p21', 'ON'))
(('CTNNB1', 'ON'), ('SMAD', 'ON'))
(('CTNNB1', 'ON'), ('VIM', 'ON'))
(('CTNNB1', 'ON'), ('p53', 'ON'))
(('CTNNB1', 'ON'), ('p63', 'ON'))
(('CTNNB1', 'ON'), ('p73', 'ON'))
(('CTNNB1', 'ON'), ('miR203', 'ON'))
(('CTNNB1', 'ON'), ('miR34', 'ON'))
(('CTNNB1', 'ON'), ('miR200', 'ON'))
(('CTNNB1', 'ON'), ('TGFbetaR', 'ON'))
(('CTNNB1', 'ON'), ('FAK', 'ON'))
(('CTNNB1', 'ON'), ('AKT1', 'OFF'))
(('CTNNB1', 'ON'), ('DKK1', 'OFF'))
(('CTNNB1', 'ON'), ('AKT2', 'OFF'))
(('CTNNB1', 'ON'), ('RAC1', 'OFF'))
(('CTNNB1', 'ON'), ('YAP1', 'OFF'))
(('CTNNB1', 'ON'), ('PIK3CA', 'OFF'))
(('CTNNB1', 'ON'), ('ERK', 'OFF'))
(('CTNNB1', 'ON'), ('p21', 'OFF'))
(('CTNNB1', 'ON'), ('SMAD', 'OFF'))
(('CTNNB1', 'ON'), ('VIM', 'OFF'))
(('CTNNB1', 'ON'), ('p53', 'OFF'))
(('CTNNB1', 'ON'), ('p63', 'OFF'))
(('CTNNB1', 'ON'), ('p73', 'OFF'))
(('CTNNB1', 'ON'), ('miR203', 'OFF'))
(('CTNNB1', 'ON'), ('miR34', 'OFF'))
(('CTNNB1', 'ON'), ('miR200', 'OFF'))
(('CTNNB1', 'ON'), ('TGFbetaR', 'OFF'))
(('CTNNB1', 'ON'), ('FAK', 'OFF'))
(('DKK1', 'ON'), ('AKT2', 'ON'))
(('DKK1', 'ON'), ('RAC1', 'ON'))
(('DKK1', 'ON'), ('YAP1', 'ON'))
(('DKK1', 'ON'), ('PIK3CA', 'ON'))
(('DKK1', 'ON'), ('ERK', 'ON'))
(('DKK1', 'ON'), ('p21', 'ON'))
(('DKK1', 'ON'), ('SMAD', 'ON'))
(('DKK1', 'ON'), ('VIM', 'ON'))
(('DKK1', 'ON'), ('p53', 'ON'))
(('DKK1', 'ON'), ('p63', 'ON'))
(('DKK1', 'ON'), ('p73', 'ON'))
(('DKK1', 'ON'), ('miR203', 'ON'))
(('DKK1', 'ON'), ('miR34', 'ON'))
(('DKK1', 'ON'), ('miR200', 'ON'))
(('DKK1', 'ON'), ('TGFbetaR', 'ON'))
(('DKK1', 'ON'), ('FAK', 'ON'))
(('DKK1', 'ON'), ('AKT1', 'OFF'))
(('DKK1', 'ON'), ('CTNNB1', 'OFF'))
(('DKK1', 'ON'), ('AKT2', 'OFF'))
(('DKK1', 'ON'), ('RAC1', 'OFF'))
(('DKK1', 'ON'), ('YAP1', 'OFF'))
(('DKK1', 'ON'), ('PIK3CA', 'OFF'))
(('DKK1', 'ON'), ('ERK', 'OFF'))
(('DKK1', 'ON'), ('p21', 'OFF'))
(('DKK1', 'ON'), ('SMAD', 'OFF'))
(('DKK1', 'ON'), ('VIM', 'OFF'))
(('DKK1', 'ON'), ('p53', 'OFF'))
(('DKK1', 'ON'), ('p63', 'OFF'))
(('DKK1', 'ON'), ('p73', 'OFF'))
(('DKK1', 'ON'), ('miR203', 'OFF'))
(('DKK1', 'ON'), ('miR34', 'OFF'))
(('DKK1', 'ON'), ('miR200', 'OFF'))
(('DKK1', 'ON'), ('TGFbetaR', 'OFF'))
(('DKK1', 'ON'), ('FAK', 'OFF'))
(('AKT2', 'ON'), ('RAC1', 'ON'))
(('AKT2', 'ON'), ('YAP1', 'ON'))
(('AKT2', 'ON'), ('PIK3CA', 'ON'))
(('AKT2', 'ON'), ('ERK', 'ON'))
(('AKT2', 'ON'), ('p21', 'ON'))
(('AKT2', 'ON'), ('SMAD', 'ON'))
(('AKT2', 'ON'), ('VIM', 'ON'))
(('AKT2', 'ON'), ('p53', 'ON'))
(('AKT2', 'ON'), ('p63', 'ON'))
(('AKT2', 'ON'), ('p73', 'ON'))
(('AKT2', 'ON'), ('miR203', 'ON'))
(('AKT2', 'ON'), ('miR34', 'ON'))
(('AKT2', 'ON'), ('miR200', 'ON'))
(('AKT2', 'ON'), ('TGFbetaR', 'ON'))
(('AKT2', 'ON'), ('FAK', 'ON'))
(('AKT2', 'ON'), ('AKT1', 'OFF'))
(('AKT2', 'ON'), ('CTNNB1', 'OFF'))
(('AKT2', 'ON'), ('DKK1', 'OFF'))
(('AKT2', 'ON'), ('RAC1', 'OFF'))
(('AKT2', 'ON'), ('YAP1', 'OFF'))
(('AKT2', 'ON'), ('PIK3CA', 'OFF'))
(('AKT2', 'ON'), ('ERK', 'OFF'))
(('AKT2', 'ON'), ('p21', 'OFF'))
(('AKT2', 'ON'), ('SMAD', 'OFF'))
(('AKT2', 'ON'), ('VIM', 'OFF'))
(('AKT2', 'ON'), ('p53', 'OFF'))
(('AKT2', 'ON'), ('p63', 'OFF'))
(('AKT2', 'ON'), ('p73', 'OFF'))
(('AKT2', 'ON'), ('miR203', 'OFF'))
(('AKT2', 'ON'), ('miR34', 'OFF'))
(('AKT2', 'ON'), ('miR200', 'OFF'))
(('AKT2', 'ON'), ('TGFbetaR', 'OFF'))
(('AKT2', 'ON'), ('FAK', 'OFF'))
(('RAC1', 'ON'), ('YAP1', 'ON'))
(('RAC1', 'ON'), ('PIK3CA', 'ON'))
(('RAC1', 'ON'), ('ERK', 'ON'))
(('RAC1', 'ON'), ('p21', 'ON'))
(('RAC1', 'ON'), ('SMAD', 'ON'))
(('RAC1', 'ON'), ('VIM', 'ON'))
(('RAC1', 'ON'), ('p53', 'ON'))
(('RAC1', 'ON'), ('p63', 'ON'))
(('RAC1', 'ON'), ('p73', 'ON'))
(('RAC1', 'ON'), ('miR203', 'ON'))
(('RAC1', 'ON'), ('miR34', 'ON'))
(('RAC1', 'ON'), ('miR200', 'ON'))
(('RAC1', 'ON'), ('TGFbetaR', 'ON'))
(('RAC1', 'ON'), ('FAK', 'ON'))
(('RAC1', 'ON'), ('AKT1', 'OFF'))
(('RAC1', 'ON'), ('CTNNB1', 'OFF'))
(('RAC1', 'ON'), ('DKK1', 'OFF'))
(('RAC1', 'ON'), ('AKT2', 'OFF'))
(('RAC1', 'ON'), ('YAP1', 'OFF'))
(('RAC1', 'ON'), ('PIK3CA', 'OFF'))
(('RAC1', 'ON'), ('ERK', 'OFF'))
(('RAC1', 'ON'), ('p21', 'OFF'))
(('RAC1', 'ON'), ('SMAD', 'OFF'))
(('RAC1', 'ON'), ('VIM', 'OFF'))
(('RAC1', 'ON'), ('p53', 'OFF'))
(('RAC1', 'ON'), ('p63', 'OFF'))
(('RAC1', 'ON'), ('p73', 'OFF'))
(('RAC1', 'ON'), ('miR203', 'OFF'))
(('RAC1', 'ON'), ('miR34', 'OFF'))
(('RAC1', 'ON'), ('miR200', 'OFF'))
(('RAC1', 'ON'), ('TGFbetaR', 'OFF'))
(('RAC1', 'ON'), ('FAK', 'OFF'))
(('YAP1', 'ON'), ('PIK3CA', 'ON'))
(('YAP1', 'ON'), ('ERK', 'ON'))
(('YAP1', 'ON'), ('p21', 'ON'))
(('YAP1', 'ON'), ('SMAD', 'ON'))
(('YAP1', 'ON'), ('VIM', 'ON'))
(('YAP1', 'ON'), ('p53', 'ON'))
(('YAP1', 'ON'), ('p63', 'ON'))
(('YAP1', 'ON'), ('p73', 'ON'))
(('YAP1', 'ON'), ('miR203', 'ON'))
(('YAP1', 'ON'), ('miR34', 'ON'))
(('YAP1', 'ON'), ('miR200', 'ON'))
(('YAP1', 'ON'), ('TGFbetaR', 'ON'))
(('YAP1', 'ON'), ('FAK', 'ON'))
(('YAP1', 'ON'), ('AKT1', 'OFF'))
(('YAP1', 'ON'), ('CTNNB1', 'OFF'))
(('YAP1', 'ON'), ('DKK1', 'OFF'))
(('YAP1', 'ON'), ('AKT2', 'OFF'))
(('YAP1', 'ON'), ('RAC1', 'OFF'))
(('YAP1', 'ON'), ('PIK3CA', 'OFF'))
(('YAP1', 'ON'), ('ERK', 'OFF'))
(('YAP1', 'ON'), ('p21', 'OFF'))
(('YAP1', 'ON'), ('SMAD', 'OFF'))
(('YAP1', 'ON'), ('VIM', 'OFF'))
(('YAP1', 'ON'), ('p53', 'OFF'))
(('YAP1', 'ON'), ('p63', 'OFF'))
(('YAP1', 'ON'), ('p73', 'OFF'))
(('YAP1', 'ON'), ('miR203', 'OFF'))
```

```
/home/marco/anaconda3/envs/MaBoSS_env/lib/python3.9/site-packages/maboss/figures.py:83: MatplotlibDeprecationWarning: normalize=None does not normalize if the sum is less than 1 but this behavior is deprecated since 3.3 until two minor releases later. After the deprecation period the default value will be normalize=True. To prevent normalization pass normalize=False 
  ax.pie(plot_line, labels=plotting_labels, radius=1.2,
```

```
(('YAP1', 'ON'), ('miR34', 'OFF'))
(('YAP1', 'ON'), ('miR200', 'OFF'))
(('YAP1', 'ON'), ('TGFbetaR', 'OFF'))
(('YAP1', 'ON'), ('FAK', 'OFF'))
(('PIK3CA', 'ON'), ('ERK', 'ON'))
(('PIK3CA', 'ON'), ('p21', 'ON'))
(('PIK3CA', 'ON'), ('SMAD', 'ON'))
(('PIK3CA', 'ON'), ('VIM', 'ON'))
(('PIK3CA', 'ON'), ('p53', 'ON'))
(('PIK3CA', 'ON'), ('p63', 'ON'))
(('PIK3CA', 'ON'), ('p73', 'ON'))
(('PIK3CA', 'ON'), ('miR203', 'ON'))
(('PIK3CA', 'ON'), ('miR34', 'ON'))
(('PIK3CA', 'ON'), ('miR200', 'ON'))
(('PIK3CA', 'ON'), ('TGFbetaR', 'ON'))
(('PIK3CA', 'ON'), ('FAK', 'ON'))
(('PIK3CA', 'ON'), ('AKT1', 'OFF'))
(('PIK3CA', 'ON'), ('CTNNB1', 'OFF'))
(('PIK3CA', 'ON'), ('DKK1', 'OFF'))
(('PIK3CA', 'ON'), ('AKT2', 'OFF'))
(('PIK3CA', 'ON'), ('RAC1', 'OFF'))
(('PIK3CA', 'ON'), ('YAP1', 'OFF'))
(('PIK3CA', 'ON'), ('ERK', 'OFF'))
(('PIK3CA', 'ON'), ('p21', 'OFF'))
(('PIK3CA', 'ON'), ('SMAD', 'OFF'))
(('PIK3CA', 'ON'), ('VIM', 'OFF'))
(('PIK3CA', 'ON'), ('p53', 'OFF'))
(('PIK3CA', 'ON'), ('p63', 'OFF'))
(('PIK3CA', 'ON'), ('p73', 'OFF'))
(('PIK3CA', 'ON'), ('miR203', 'OFF'))
(('PIK3CA', 'ON'), ('miR34', 'OFF'))
(('PIK3CA', 'ON'), ('miR200', 'OFF'))
(('PIK3CA', 'ON'), ('TGFbetaR', 'OFF'))
(('PIK3CA', 'ON'), ('FAK', 'OFF'))
(('ERK', 'ON'), ('p21', 'ON'))
(('ERK', 'ON'), ('SMAD', 'ON'))
(('ERK', 'ON'), ('VIM', 'ON'))
(('ERK', 'ON'), ('p53', 'ON'))
(('ERK', 'ON'), ('p63', 'ON'))
(('ERK', 'ON'), ('p73', 'ON'))
(('ERK', 'ON'), ('miR203', 'ON'))
(('ERK', 'ON'), ('miR34', 'ON'))
(('ERK', 'ON'), ('miR200', 'ON'))
(('ERK', 'ON'), ('TGFbetaR', 'ON'))
(('ERK', 'ON'), ('FAK', 'ON'))
(('ERK', 'ON'), ('AKT1', 'OFF'))
(('ERK', 'ON'), ('CTNNB1', 'OFF'))
(('ERK', 'ON'), ('DKK1', 'OFF'))
(('ERK', 'ON'), ('AKT2', 'OFF'))
(('ERK', 'ON'), ('RAC1', 'OFF'))
(('ERK', 'ON'), ('YAP1', 'OFF'))
(('ERK', 'ON'), ('PIK3CA', 'OFF'))
(('ERK', 'ON'), ('p21', 'OFF'))
(('ERK', 'ON'), ('SMAD', 'OFF'))
(('ERK', 'ON'), ('VIM', 'OFF'))
(('ERK', 'ON'), ('p53', 'OFF'))
(('ERK', 'ON'), ('p63', 'OFF'))
(('ERK', 'ON'), ('p73', 'OFF'))
(('ERK', 'ON'), ('miR203', 'OFF'))
(('ERK', 'ON'), ('miR34', 'OFF'))
(('ERK', 'ON'), ('miR200', 'OFF'))
(('ERK', 'ON'), ('TGFbetaR', 'OFF'))
(('ERK', 'ON'), ('FAK', 'OFF'))
(('p21', 'ON'), ('SMAD', 'ON'))
(('p21', 'ON'), ('VIM', 'ON'))
(('p21', 'ON'), ('p53', 'ON'))
(('p21', 'ON'), ('p63', 'ON'))
(('p21', 'ON'), ('p73', 'ON'))
(('p21', 'ON'), ('miR203', 'ON'))
(('p21', 'ON'), ('miR34', 'ON'))
(('p21', 'ON'), ('miR200', 'ON'))
(('p21', 'ON'), ('TGFbetaR', 'ON'))
(('p21', 'ON'), ('FAK', 'ON'))
(('p21', 'ON'), ('AKT1', 'OFF'))
(('p21', 'ON'), ('CTNNB1', 'OFF'))
(('p21', 'ON'), ('DKK1', 'OFF'))
(('p21', 'ON'), ('AKT2', 'OFF'))
(('p21', 'ON'), ('RAC1', 'OFF'))
(('p21', 'ON'), ('YAP1', 'OFF'))
(('p21', 'ON'), ('PIK3CA', 'OFF'))
(('p21', 'ON'), ('ERK', 'OFF'))
(('p21', 'ON'), ('SMAD', 'OFF'))
(('p21', 'ON'), ('VIM', 'OFF'))
(('p21', 'ON'), ('p53', 'OFF'))
(('p21', 'ON'), ('p63', 'OFF'))
(('p21', 'ON'), ('p73', 'OFF'))
(('p21', 'ON'), ('miR203', 'OFF'))
(('p21', 'ON'), ('miR34', 'OFF'))
(('p21', 'ON'), ('miR200', 'OFF'))
(('p21', 'ON'), ('TGFbetaR', 'OFF'))
(('p21', 'ON'), ('FAK', 'OFF'))
(('SMAD', 'ON'), ('VIM', 'ON'))
(('SMAD', 'ON'), ('p53', 'ON'))
(('SMAD', 'ON'), ('p63', 'ON'))
(('SMAD', 'ON'), ('p73', 'ON'))
(('SMAD', 'ON'), ('miR203', 'ON'))
(('SMAD', 'ON'), ('miR34', 'ON'))
(('SMAD', 'ON'), ('miR200', 'ON'))
(('SMAD', 'ON'), ('TGFbetaR', 'ON'))
(('SMAD', 'ON'), ('FAK', 'ON'))
(('SMAD', 'ON'), ('AKT1', 'OFF'))
(('SMAD', 'ON'), ('CTNNB1', 'OFF'))
(('SMAD', 'ON'), ('DKK1', 'OFF'))
(('SMAD', 'ON'), ('AKT2', 'OFF'))
(('SMAD', 'ON'), ('RAC1', 'OFF'))
(('SMAD', 'ON'), ('YAP1', 'OFF'))
(('SMAD', 'ON'), ('PIK3CA', 'OFF'))
(('SMAD', 'ON'), ('ERK', 'OFF'))
(('SMAD', 'ON'), ('p21', 'OFF'))
(('SMAD', 'ON'), ('VIM', 'OFF'))
(('SMAD', 'ON'), ('p53', 'OFF'))
(('SMAD', 'ON'), ('p63', 'OFF'))
(('SMAD', 'ON'), ('p73', 'OFF'))
(('SMAD', 'ON'), ('miR203', 'OFF'))
(('SMAD', 'ON'), ('miR34', 'OFF'))
(('SMAD', 'ON'), ('miR200', 'OFF'))
(('SMAD', 'ON'), ('TGFbetaR', 'OFF'))
(('SMAD', 'ON'), ('FAK', 'OFF'))
(('VIM', 'ON'), ('p53', 'ON'))
(('VIM', 'ON'), ('p63', 'ON'))
(('VIM', 'ON'), ('p73', 'ON'))
(('VIM', 'ON'), ('miR203', 'ON'))
(('VIM', 'ON'), ('miR34', 'ON'))
(('VIM', 'ON'), ('miR200', 'ON'))
(('VIM', 'ON'), ('TGFbetaR', 'ON'))
(('VIM', 'ON'), ('FAK', 'ON'))
(('VIM', 'ON'), ('AKT1', 'OFF'))
(('VIM', 'ON'), ('CTNNB1', 'OFF'))
(('VIM', 'ON'), ('DKK1', 'OFF'))
(('VIM', 'ON'), ('AKT2', 'OFF'))
(('VIM', 'ON'), ('RAC1', 'OFF'))
(('VIM', 'ON'), ('YAP1', 'OFF'))
(('VIM', 'ON'), ('PIK3CA', 'OFF'))
(('VIM', 'ON'), ('ERK', 'OFF'))
(('VIM', 'ON'), ('p21', 'OFF'))
(('VIM', 'ON'), ('SMAD', 'OFF'))
(('VIM', 'ON'), ('p53', 'OFF'))
(('VIM', 'ON'), ('p63', 'OFF'))
(('VIM', 'ON'), ('p73', 'OFF'))
(('VIM', 'ON'), ('miR203', 'OFF'))
(('VIM', 'ON'), ('miR34', 'OFF'))
(('VIM', 'ON'), ('miR200', 'OFF'))
(('VIM', 'ON'), ('TGFbetaR', 'OFF'))
(('VIM', 'ON'), ('FAK', 'OFF'))
(('p53', 'ON'), ('p63', 'ON'))
(('p53', 'ON'), ('p73', 'ON'))
(('p53', 'ON'), ('miR203', 'ON'))
(('p53', 'ON'), ('miR34', 'ON'))
(('p53', 'ON'), ('miR200', 'ON'))
(('p53', 'ON'), ('TGFbetaR', 'ON'))
(('p53', 'ON'), ('FAK', 'ON'))
(('p53', 'ON'), ('AKT1', 'OFF'))
(('p53', 'ON'), ('CTNNB1', 'OFF'))
(('p53', 'ON'), ('DKK1', 'OFF'))
(('p53', 'ON'), ('AKT2', 'OFF'))
(('p53', 'ON'), ('RAC1', 'OFF'))
(('p53', 'ON'), ('YAP1', 'OFF'))
(('p53', 'ON'), ('PIK3CA', 'OFF'))
(('p53', 'ON'), ('ERK', 'OFF'))
(('p53', 'ON'), ('p21', 'OFF'))
(('p53', 'ON'), ('SMAD', 'OFF'))
(('p53', 'ON'), ('VIM', 'OFF'))
(('p53', 'ON'), ('p63', 'OFF'))
(('p53', 'ON'), ('p73', 'OFF'))
(('p53', 'ON'), ('miR203', 'OFF'))
(('p53', 'ON'), ('miR34', 'OFF'))
(('p53', 'ON'), ('miR200', 'OFF'))
(('p53', 'ON'), ('TGFbetaR', 'OFF'))
(('p53', 'ON'), ('FAK', 'OFF'))
(('p63', 'ON'), ('p73', 'ON'))
(('p63', 'ON'), ('miR203', 'ON'))
(('p63', 'ON'), ('miR34', 'ON'))
(('p63', 'ON'), ('miR200', 'ON'))
(('p63', 'ON'), ('TGFbetaR', 'ON'))
(('p63', 'ON'), ('FAK', 'ON'))
(('p63', 'ON'), ('AKT1', 'OFF'))
(('p63', 'ON'), ('CTNNB1', 'OFF'))
(('p63', 'ON'), ('DKK1', 'OFF'))
(('p63', 'ON'), ('AKT2', 'OFF'))
(('p63', 'ON'), ('RAC1', 'OFF'))
(('p63', 'ON'), ('YAP1', 'OFF'))
(('p63', 'ON'), ('PIK3CA', 'OFF'))
(('p63', 'ON'), ('ERK', 'OFF'))
(('p63', 'ON'), ('p21', 'OFF'))
(('p63', 'ON'), ('SMAD', 'OFF'))
(('p63', 'ON'), ('VIM', 'OFF'))
(('p63', 'ON'), ('p53', 'OFF'))
(('p63', 'ON'), ('p73', 'OFF'))
(('p63', 'ON'), ('miR203', 'OFF'))
(('p63', 'ON'), ('miR34', 'OFF'))
(('p63', 'ON'), ('miR200', 'OFF'))
(('p63', 'ON'), ('TGFbetaR', 'OFF'))
(('p63', 'ON'), ('FAK', 'OFF'))
(('p73', 'ON'), ('miR203', 'ON'))
(('p73', 'ON'), ('miR34', 'ON'))
(('p73', 'ON'), ('miR200', 'ON'))
(('p73', 'ON'), ('TGFbetaR', 'ON'))
(('p73', 'ON'), ('FAK', 'ON'))
(('p73', 'ON'), ('AKT1', 'OFF'))
(('p73', 'ON'), ('CTNNB1', 'OFF'))
(('p73', 'ON'), ('DKK1', 'OFF'))
(('p73', 'ON'), ('AKT2', 'OFF'))
(('p73', 'ON'), ('RAC1', 'OFF'))
(('p73', 'ON'), ('YAP1', 'OFF'))
(('p73', 'ON'), ('PIK3CA', 'OFF'))
(('p73', 'ON'), ('ERK', 'OFF'))
(('p73', 'ON'), ('p21', 'OFF'))
(('p73', 'ON'), ('SMAD', 'OFF'))
(('p73', 'ON'), ('VIM', 'OFF'))
(('p73', 'ON'), ('p53', 'OFF'))
(('p73', 'ON'), ('p63', 'OFF'))
(('p73', 'ON'), ('miR203', 'OFF'))
(('p73', 'ON'), ('miR34', 'OFF'))
(('p73', 'ON'), ('miR200', 'OFF'))
(('p73', 'ON'), ('TGFbetaR', 'OFF'))
(('p73', 'ON'), ('FAK', 'OFF'))
(('miR203', 'ON'), ('miR34', 'ON'))
(('miR203', 'ON'), ('miR200', 'ON'))
(('miR203', 'ON'), ('TGFbetaR', 'ON'))
(('miR203', 'ON'), ('FAK', 'ON'))
(('miR203', 'ON'), ('AKT1', 'OFF'))
(('miR203', 'ON'), ('CTNNB1', 'OFF'))
(('miR203', 'ON'), ('DKK1', 'OFF'))
(('miR203', 'ON'), ('AKT2', 'OFF'))
(('miR203', 'ON'), ('RAC1', 'OFF'))
(('miR203', 'ON'), ('YAP1', 'OFF'))
(('miR203', 'ON'), ('PIK3CA', 'OFF'))
(('miR203', 'ON'), ('ERK', 'OFF'))
(('miR203', 'ON'), ('p21', 'OFF'))
(('miR203', 'ON'), ('SMAD', 'OFF'))
(('miR203', 'ON'), ('VIM', 'OFF'))
(('miR203', 'ON'), ('p53', 'OFF'))
(('miR203', 'ON'), ('p63', 'OFF'))
(('miR203', 'ON'), ('p73', 'OFF'))
(('miR203', 'ON'), ('miR34', 'OFF'))
(('miR203', 'ON'), ('miR200', 'OFF'))
(('miR203', 'ON'), ('TGFbetaR', 'OFF'))
(('miR203', 'ON'), ('FAK', 'OFF'))
(('miR34', 'ON'), ('miR200', 'ON'))
(('miR34', 'ON'), ('TGFbetaR', 'ON'))
(('miR34', 'ON'), ('FAK', 'ON'))
(('miR34', 'ON'), ('AKT1', 'OFF'))
(('miR34', 'ON'), ('CTNNB1', 'OFF'))
(('miR34', 'ON'), ('DKK1', 'OFF'))
(('miR34', 'ON'), ('AKT2', 'OFF'))
(('miR34', 'ON'), ('RAC1', 'OFF'))
(('miR34', 'ON'), ('YAP1', 'OFF'))
(('miR34', 'ON'), ('PIK3CA', 'OFF'))
(('miR34', 'ON'), ('ERK', 'OFF'))
(('miR34', 'ON'), ('p21', 'OFF'))
(('miR34', 'ON'), ('SMAD', 'OFF'))
(('miR34', 'ON'), ('VIM', 'OFF'))
(('miR34', 'ON'), ('p53', 'OFF'))
(('miR34', 'ON'), ('p63', 'OFF'))
(('miR34', 'ON'), ('p73', 'OFF'))
(('miR34', 'ON'), ('miR203', 'OFF'))
(('miR34', 'ON'), ('miR200', 'OFF'))
(('miR34', 'ON'), ('TGFbetaR', 'OFF'))
(('miR34', 'ON'), ('FAK', 'OFF'))
(('miR200', 'ON'), ('TGFbetaR', 'ON'))
(('miR200', 'ON'), ('FAK', 'ON'))
(('miR200', 'ON'), ('AKT1', 'OFF'))
(('miR200', 'ON'), ('CTNNB1', 'OFF'))
(('miR200', 'ON'), ('DKK1', 'OFF'))
(('miR200', 'ON'), ('AKT2', 'OFF'))
(('miR200', 'ON'), ('RAC1', 'OFF'))
(('miR200', 'ON'), ('YAP1', 'OFF'))
(('miR200', 'ON'), ('PIK3CA', 'OFF'))
(('miR200', 'ON'), ('ERK', 'OFF'))
(('miR200', 'ON'), ('p21', 'OFF'))
(('miR200', 'ON'), ('SMAD', 'OFF'))
(('miR200', 'ON'), ('VIM', 'OFF'))
(('miR200', 'ON'), ('p53', 'OFF'))
(('miR200', 'ON'), ('p63', 'OFF'))
(('miR200', 'ON'), ('p73', 'OFF'))
(('miR200', 'ON'), ('miR203', 'OFF'))
(('miR200', 'ON'), ('miR34', 'OFF'))
(('miR200', 'ON'), ('TGFbetaR', 'OFF'))
(('miR200', 'ON'), ('FAK', 'OFF'))
(('TGFbetaR', 'ON'), ('FAK', 'ON'))
(('TGFbetaR', 'ON'), ('AKT1', 'OFF'))
(('TGFbetaR', 'ON'), ('CTNNB1', 'OFF'))
(('TGFbetaR', 'ON'), ('DKK1', 'OFF'))
(('TGFbetaR', 'ON'), ('AKT2', 'OFF'))
(('TGFbetaR', 'ON'), ('RAC1', 'OFF'))
(('TGFbetaR', 'ON'), ('YAP1', 'OFF'))
(('TGFbetaR', 'ON'), ('PIK3CA', 'OFF'))
(('TGFbetaR', 'ON'), ('ERK', 'OFF'))
(('TGFbetaR', 'ON'), ('p21', 'OFF'))
(('TGFbetaR', 'ON'), ('SMAD', 'OFF'))
(('TGFbetaR', 'ON'), ('VIM', 'OFF'))
(('TGFbetaR', 'ON'), ('p53', 'OFF'))
(('TGFbetaR', 'ON'), ('p63', 'OFF'))
(('TGFbetaR', 'ON'), ('p73', 'OFF'))
(('TGFbetaR', 'ON'), ('miR203', 'OFF'))
(('TGFbetaR', 'ON'), ('miR34', 'OFF'))
(('TGFbetaR', 'ON'), ('miR200', 'OFF'))
(('TGFbetaR', 'ON'), ('FAK', 'OFF'))
(('FAK', 'ON'), ('AKT1', 'OFF'))
(('FAK', 'ON'), ('CTNNB1', 'OFF'))
(('FAK', 'ON'), ('DKK1', 'OFF'))
(('FAK', 'ON'), ('AKT2', 'OFF'))
(('FAK', 'ON'), ('RAC1', 'OFF'))
(('FAK', 'ON'), ('YAP1', 'OFF'))
(('FAK', 'ON'), ('PIK3CA', 'OFF'))
(('FAK', 'ON'), ('ERK', 'OFF'))
(('FAK', 'ON'), ('p21', 'OFF'))
(('FAK', 'ON'), ('SMAD', 'OFF'))
(('FAK', 'ON'), ('VIM', 'OFF'))
(('FAK', 'ON'), ('p53', 'OFF'))
(('FAK', 'ON'), ('p63', 'OFF'))
(('FAK', 'ON'), ('p73', 'OFF'))
(('FAK', 'ON'), ('miR203', 'OFF'))
(('FAK', 'ON'), ('miR34', 'OFF'))
(('FAK', 'ON'), ('miR200', 'OFF'))
(('FAK', 'ON'), ('TGFbetaR', 'OFF'))
(('AKT1', 'OFF'), ('CTNNB1', 'OFF'))
(('AKT1', 'OFF'), ('DKK1', 'OFF'))
(('AKT1', 'OFF'), ('AKT2', 'OFF'))
(('AKT1', 'OFF'), ('RAC1', 'OFF'))
(('AKT1', 'OFF'), ('YAP1', 'OFF'))
(('AKT1', 'OFF'), ('PIK3CA', 'OFF'))
(('AKT1', 'OFF'), ('ERK', 'OFF'))
(('AKT1', 'OFF'), ('p21', 'OFF'))
(('AKT1', 'OFF'), ('SMAD', 'OFF'))
(('AKT1', 'OFF'), ('VIM', 'OFF'))
(('AKT1', 'OFF'), ('p53', 'OFF'))
(('AKT1', 'OFF'), ('p63', 'OFF'))
(('AKT1', 'OFF'), ('p73', 'OFF'))
(('AKT1', 'OFF'), ('miR203', 'OFF'))
(('AKT1', 'OFF'), ('miR34', 'OFF'))
(('AKT1', 'OFF'), ('miR200', 'OFF'))
(('AKT1', 'OFF'), ('TGFbetaR', 'OFF'))
(('AKT1', 'OFF'), ('FAK', 'OFF'))
(('CTNNB1', 'OFF'), ('DKK1', 'OFF'))
(('CTNNB1', 'OFF'), ('AKT2', 'OFF'))
(('CTNNB1', 'OFF'), ('RAC1', 'OFF'))
(('CTNNB1', 'OFF'), ('YAP1', 'OFF'))
(('CTNNB1', 'OFF'), ('PIK3CA', 'OFF'))
(('CTNNB1', 'OFF'), ('ERK', 'OFF'))
(('CTNNB1', 'OFF'), ('p21', 'OFF'))
(('CTNNB1', 'OFF'), ('SMAD', 'OFF'))
(('CTNNB1', 'OFF'), ('VIM', 'OFF'))
(('CTNNB1', 'OFF'), ('p53', 'OFF'))
(('CTNNB1', 'OFF'), ('p63', 'OFF'))
(('CTNNB1', 'OFF'), ('p73', 'OFF'))
(('CTNNB1', 'OFF'), ('miR203', 'OFF'))
(('CTNNB1', 'OFF'), ('miR34', 'OFF'))
(('CTNNB1', 'OFF'), ('miR200', 'OFF'))
(('CTNNB1', 'OFF'), ('TGFbetaR', 'OFF'))
(('CTNNB1', 'OFF'), ('FAK', 'OFF'))
(('DKK1', 'OFF'), ('AKT2', 'OFF'))
(('DKK1', 'OFF'), ('RAC1', 'OFF'))
(('DKK1', 'OFF'), ('YAP1', 'OFF'))
(('DKK1', 'OFF'), ('PIK3CA', 'OFF'))
(('DKK1', 'OFF'), ('ERK', 'OFF'))
(('DKK1', 'OFF'), ('p21', 'OFF'))
(('DKK1', 'OFF'), ('SMAD', 'OFF'))
(('DKK1', 'OFF'), ('VIM', 'OFF'))
(('DKK1', 'OFF'), ('p53', 'OFF'))
(('DKK1', 'OFF'), ('p63', 'OFF'))
(('DKK1', 'OFF'), ('p73', 'OFF'))
(('DKK1', 'OFF'), ('miR203', 'OFF'))
(('DKK1', 'OFF'), ('miR34', 'OFF'))
(('DKK1', 'OFF'), ('miR200', 'OFF'))
(('DKK1', 'OFF'), ('TGFbetaR', 'OFF'))
(('DKK1', 'OFF'), ('FAK', 'OFF'))
(('AKT2', 'OFF'), ('RAC1', 'OFF'))
(('AKT2', 'OFF'), ('YAP1', 'OFF'))
(('AKT2', 'OFF'), ('PIK3CA', 'OFF'))
(('AKT2', 'OFF'), ('ERK', 'OFF'))
(('AKT2', 'OFF'), ('p21', 'OFF'))
(('AKT2', 'OFF'), ('SMAD', 'OFF'))
(('AKT2', 'OFF'), ('VIM', 'OFF'))
(('AKT2', 'OFF'), ('p53', 'OFF'))
(('AKT2', 'OFF'), ('p63', 'OFF'))
(('AKT2', 'OFF'), ('p73', 'OFF'))
(('AKT2', 'OFF'), ('miR203', 'OFF'))
(('AKT2', 'OFF'), ('miR34', 'OFF'))
(('AKT2', 'OFF'), ('miR200', 'OFF'))
(('AKT2', 'OFF'), ('TGFbetaR', 'OFF'))
(('AKT2', 'OFF'), ('FAK', 'OFF'))
(('RAC1', 'OFF'), ('YAP1', 'OFF'))
(('RAC1', 'OFF'), ('PIK3CA', 'OFF'))
(('RAC1', 'OFF'), ('ERK', 'OFF'))
(('RAC1', 'OFF'), ('p21', 'OFF'))
(('RAC1', 'OFF'), ('SMAD', 'OFF'))
(('RAC1', 'OFF'), ('VIM', 'OFF'))
(('RAC1', 'OFF'), ('p53', 'OFF'))
(('RAC1', 'OFF'), ('p63', 'OFF'))
(('RAC1', 'OFF'), ('p73', 'OFF'))
(('RAC1', 'OFF'), ('miR203', 'OFF'))
(('RAC1', 'OFF'), ('miR34', 'OFF'))
(('RAC1', 'OFF'), ('miR200', 'OFF'))
(('RAC1', 'OFF'), ('TGFbetaR', 'OFF'))
(('RAC1', 'OFF'), ('FAK', 'OFF'))
(('YAP1', 'OFF'), ('PIK3CA', 'OFF'))
(('YAP1', 'OFF'), ('ERK', 'OFF'))
(('YAP1', 'OFF'), ('p21', 'OFF'))
(('YAP1', 'OFF'), ('SMAD', 'OFF'))
(('YAP1', 'OFF'), ('VIM', 'OFF'))
(('YAP1', 'OFF'), ('p53', 'OFF'))
(('YAP1', 'OFF'), ('p63', 'OFF'))
(('YAP1', 'OFF'), ('p73', 'OFF'))
(('YAP1', 'OFF'), ('miR203', 'OFF'))
(('YAP1', 'OFF'), ('miR34', 'OFF'))
(('YAP1', 'OFF'), ('miR200', 'OFF'))
(('YAP1', 'OFF'), ('TGFbetaR', 'OFF'))
(('YAP1', 'OFF'), ('FAK', 'OFF'))
(('PIK3CA', 'OFF'), ('ERK', 'OFF'))
(('PIK3CA', 'OFF'), ('p21', 'OFF'))
(('PIK3CA', 'OFF'), ('SMAD', 'OFF'))
(('PIK3CA', 'OFF'), ('VIM', 'OFF'))
(('PIK3CA', 'OFF'), ('p53', 'OFF'))
(('PIK3CA', 'OFF'), ('p63', 'OFF'))
(('PIK3CA', 'OFF'), ('p73', 'OFF'))
(('PIK3CA', 'OFF'), ('miR203', 'OFF'))
(('PIK3CA', 'OFF'), ('miR34', 'OFF'))
(('PIK3CA', 'OFF'), ('miR200', 'OFF'))
(('PIK3CA', 'OFF'), ('TGFbetaR', 'OFF'))
(('PIK3CA', 'OFF'), ('FAK', 'OFF'))
(('ERK', 'OFF'), ('p21', 'OFF'))
(('ERK', 'OFF'), ('SMAD', 'OFF'))
(('ERK', 'OFF'), ('VIM', 'OFF'))
(('ERK', 'OFF'), ('p53', 'OFF'))
(('ERK', 'OFF'), ('p63', 'OFF'))
(('ERK', 'OFF'), ('p73', 'OFF'))
(('ERK', 'OFF'), ('miR203', 'OFF'))
(('ERK', 'OFF'), ('miR34', 'OFF'))
(('ERK', 'OFF'), ('miR200', 'OFF'))
(('ERK', 'OFF'), ('TGFbetaR', 'OFF'))
(('ERK', 'OFF'), ('FAK', 'OFF'))
(('p21', 'OFF'), ('SMAD', 'OFF'))
(('p21', 'OFF'), ('VIM', 'OFF'))
(('p21', 'OFF'), ('p53', 'OFF'))
(('p21', 'OFF'), ('p63', 'OFF'))
(('p21', 'OFF'), ('p73', 'OFF'))
(('p21', 'OFF'), ('miR203', 'OFF'))
(('p21', 'OFF'), ('miR34', 'OFF'))
(('p21', 'OFF'), ('miR200', 'OFF'))
(('p21', 'OFF'), ('TGFbetaR', 'OFF'))
(('p21', 'OFF'), ('FAK', 'OFF'))
(('SMAD', 'OFF'), ('VIM', 'OFF'))
(('SMAD', 'OFF'), ('p53', 'OFF'))
(('SMAD', 'OFF'), ('p63', 'OFF'))
(('SMAD', 'OFF'), ('p73', 'OFF'))
(('SMAD', 'OFF'), ('miR203', 'OFF'))
(('SMAD', 'OFF'), ('miR34', 'OFF'))
(('SMAD', 'OFF'), ('miR200', 'OFF'))
(('SMAD', 'OFF'), ('TGFbetaR', 'OFF'))
(('SMAD', 'OFF'), ('FAK', 'OFF'))
(('VIM', 'OFF'), ('p53', 'OFF'))
(('VIM', 'OFF'), ('p63', 'OFF'))
(('VIM', 'OFF'), ('p73', 'OFF'))
(('VIM', 'OFF'), ('miR203', 'OFF'))
(('VIM', 'OFF'), ('miR34', 'OFF'))
(('VIM', 'OFF'), ('miR200', 'OFF'))
(('VIM', 'OFF'), ('TGFbetaR', 'OFF'))
(('VIM', 'OFF'), ('FAK', 'OFF'))
(('p53', 'OFF'), ('p63', 'OFF'))
(('p53', 'OFF'), ('p73', 'OFF'))
(('p53', 'OFF'), ('miR203', 'OFF'))
(('p53', 'OFF'), ('miR34', 'OFF'))
(('p53', 'OFF'), ('miR200', 'OFF'))
(('p53', 'OFF'), ('TGFbetaR', 'OFF'))
(('p53', 'OFF'), ('FAK', 'OFF'))
(('p63', 'OFF'), ('p73', 'OFF'))
(('p63', 'OFF'), ('miR203', 'OFF'))
(('p63', 'OFF'), ('miR34', 'OFF'))
(('p63', 'OFF'), ('miR200', 'OFF'))
(('p63', 'OFF'), ('TGFbetaR', 'OFF'))
(('p63', 'OFF'), ('FAK', 'OFF'))
(('p73', 'OFF'), ('miR203', 'OFF'))
(('p73', 'OFF'), ('miR34', 'OFF'))
(('p73', 'OFF'), ('miR200', 'OFF'))
(('p73', 'OFF'), ('TGFbetaR', 'OFF'))
(('p73', 'OFF'), ('FAK', 'OFF'))
(('miR203', 'OFF'), ('miR34', 'OFF'))
(('miR203', 'OFF'), ('miR200', 'OFF'))
(('miR203', 'OFF'), ('TGFbetaR', 'OFF'))
(('miR203', 'OFF'), ('FAK', 'OFF'))
(('miR34', 'OFF'), ('miR200', 'OFF'))
(('miR34', 'OFF'), ('TGFbetaR', 'OFF'))
(('miR34', 'OFF'), ('FAK', 'OFF'))
(('miR200', 'OFF'), ('TGFbetaR', 'OFF'))
(('miR200', 'OFF'), ('FAK', 'OFF'))
(('TGFbetaR', 'OFF'), ('FAK', 'OFF'))
```

In [68]:

```
filtered_res = filter_sensititivy(res_single, node="ECM_degrad", maximum=0)
print(filtered_res.keys())
```

```
dict_keys([('AKT1', 'ON'), ('CTNNB1', 'ON'), ('DKK1', 'ON'), ('AKT2', 'ON'), ('RAC1', 'ON'), ('YAP1', 'ON'), ('PIK3CA', 'ON'), ('ERK', 'ON'), ('p21', 'ON'), ('SMAD', 'ON'), ('VIM', 'ON'), ('p53', 'ON'), ('p73', 'ON'), ('miR203', 'ON'), ('miR34', 'ON'), ('miR200', 'ON'), ('TGFbetaR', 'ON'), ('FAK', 'ON'), ('AKT1', 'OFF'), ('CTNNB1', 'OFF'), ('DKK1', 'OFF'), ('AKT2', 'OFF'), ('RAC1', 'OFF'), ('YAP1', 'OFF'), ('PIK3CA', 'OFF'), ('ERK', 'OFF'), ('p21', 'OFF'), ('SMAD', 'OFF'), ('VIM', 'OFF'), ('p53', 'OFF'), ('p63', 'OFF'), ('p73', 'OFF'), ('miR203', 'OFF'), ('miR34', 'OFF'), ('miR200', 'OFF'), ('TGFbetaR', 'OFF'), ('FAK', 'OFF')])
```

In [69]:

```
filtered_res1 = filter_sensititivy(res_single, node="EMT", maximum=0)
print(filtered_res1.keys())
```

```
dict_keys([('p53', 'ON')])
```

In [70]:

```
filtered_res2 = filter_sensititivy(res_single, node="Migration", maximum=0)
print(filtered_res2.keys())
```

```
dict_keys([('AKT1', 'ON'), ('CTNNB1', 'ON'), ('p53', 'ON'), ('p63', 'ON'), ('p73', 'ON'), ('miR203', 'ON'), ('miR34', 'ON'), ('miR200', 'ON'), ('AKT2', 'OFF'), ('ERK', 'OFF'), ('SMAD', 'OFF'), ('VIM', 'OFF')])
```

In [71]:

```
for elem in filtered_res:
    for elem2 in filtered_res1:
        for elem3 in filtered_res2:
            if elem == elem2 and elem2 == elem3:
                print(elem)
```

```
('p53', 'ON')
```

Conclusion: among all of the possible mutations, in the perfect condition for proliferation, the common candidates to stop both MMPs release, Migration and EMT phenotypes are p53, p73 and miR34. Those are possible gene that can be overexpressed within the PhysiBoSS simulation, to see if effectively they block the invasion process.

In [72]:

```
filtered_res = filter_sensititivy(res_double, node="EMT", maximum=0)
print(filtered_res.keys())
```

```
dict_keys([(('AKT1', 'ON'), ('p53', 'ON')), (('DKK1', 'ON'), ('p53', 'ON')), (('AKT2', 'ON'), ('p53', 'ON')), (('RAC1', 'ON'), ('p53', 'ON')), (('YAP1', 'ON'), ('p53', 'ON')), (('PIK3CA', 'ON'), ('p53', 'ON')), (('ERK', 'ON'), ('p53', 'ON')), (('p21', 'ON'), ('p53', 'ON')), (('SMAD', 'ON'), ('p53', 'ON')), (('VIM', 'ON'), ('p53', 'ON')), (('p53', 'ON'), ('p63', 'ON')), (('p53', 'ON'), ('p73', 'ON')), (('p53', 'ON'), ('miR203', 'ON')), (('p53', 'ON'), ('miR34', 'ON')), (('p53', 'ON'), ('miR200', 'ON')), (('p53', 'ON'), ('TGFbetaR', 'ON')), (('p53', 'ON'), ('AKT1', 'OFF')), (('p53', 'ON'), ('CTNNB1', 'OFF')), (('p53', 'ON'), ('DKK1', 'OFF')), (('p53', 'ON'), ('AKT2', 'OFF')), (('p53', 'ON'), ('RAC1', 'OFF')), (('p53', 'ON'), ('YAP1', 'OFF')), (('p53', 'ON'), ('PIK3CA', 'OFF')), (('p53', 'ON'), ('ERK', 'OFF')), (('p53', 'ON'), ('p21', 'OFF')), (('p53', 'ON'), ('SMAD', 'OFF')), (('p53', 'ON'), ('VIM', 'OFF')), (('p53', 'ON'), ('p63', 'OFF')), (('p53', 'ON'), ('p73', 'OFF')), (('p53', 'ON'), ('miR203', 'OFF')), (('p53', 'ON'), ('miR34', 'OFF')), (('p53', 'ON'), ('miR200', 'OFF')), (('p53', 'ON'), ('TGFbetaR', 'OFF')), (('p53', 'ON'), ('FAK', 'OFF')), (('p63', 'ON'), ('FAK', 'OFF')), (('p73', 'ON'), ('FAK', 'OFF')), (('miR34', 'ON'), ('FAK', 'OFF')), (('miR200', 'ON'), ('FAK', 'OFF'))])
```

In [73]:

```
filtered_res1 = filter_sensititivy(res_double, node="ECM_degrad", maximum=0)
print(filtered_res.keys())
```

```
dict_keys([(('AKT1', 'ON'), ('p53', 'ON')), (('DKK1', 'ON'), ('p53', 'ON')), (('AKT2', 'ON'), ('p53', 'ON')), (('RAC1', 'ON'), ('p53', 'ON')), (('YAP1', 'ON'), ('p53', 'ON')), (('PIK3CA', 'ON'), ('p53', 'ON')), (('ERK', 'ON'), ('p53', 'ON')), (('p21', 'ON'), ('p53', 'ON')), (('SMAD', 'ON'), ('p53', 'ON')), (('VIM', 'ON'), ('p53', 'ON')), (('p53', 'ON'), ('p63', 'ON')), (('p53', 'ON'), ('p73', 'ON')), (('p53', 'ON'), ('miR203', 'ON')), (('p53', 'ON'), ('miR34', 'ON')), (('p53', 'ON'), ('miR200', 'ON')), (('p53', 'ON'), ('TGFbetaR', 'ON')), (('p53', 'ON'), ('AKT1', 'OFF')), (('p53', 'ON'), ('CTNNB1', 'OFF')), (('p53', 'ON'), ('DKK1', 'OFF')), (('p53', 'ON'), ('AKT2', 'OFF')), (('p53', 'ON'), ('RAC1', 'OFF')), (('p53', 'ON'), ('YAP1', 'OFF')), (('p53', 'ON'), ('PIK3CA', 'OFF')), (('p53', 'ON'), ('ERK', 'OFF')), (('p53', 'ON'), ('p21', 'OFF')), (('p53', 'ON'), ('SMAD', 'OFF')), (('p53', 'ON'), ('VIM', 'OFF')), (('p53', 'ON'), ('p63', 'OFF')), (('p53', 'ON'), ('p73', 'OFF')), (('p53', 'ON'), ('miR203', 'OFF')), (('p53', 'ON'), ('miR34', 'OFF')), (('p53', 'ON'), ('miR200', 'OFF')), (('p53', 'ON'), ('TGFbetaR', 'OFF')), (('p53', 'ON'), ('FAK', 'OFF')), (('p63', 'ON'), ('FAK', 'OFF')), (('p73', 'ON'), ('FAK', 'OFF')), (('miR34', 'ON'), ('FAK', 'OFF')), (('miR200', 'ON'), ('FAK', 'OFF'))])
```

In [74]:

```
filtered_res2 = filter_sensititivy(res_double, node="Migration", maximum=0)
print(filtered_res.keys())
```

```
dict_keys([(('AKT1', 'ON'), ('p53', 'ON')), (('DKK1', 'ON'), ('p53', 'ON')), (('AKT2', 'ON'), ('p53', 'ON')), (('RAC1', 'ON'), ('p53', 'ON')), (('YAP1', 'ON'), ('p53', 'ON')), (('PIK3CA', 'ON'), ('p53', 'ON')), (('ERK', 'ON'), ('p53', 'ON')), (('p21', 'ON'), ('p53', 'ON')), (('SMAD', 'ON'), ('p53', 'ON')), (('VIM', 'ON'), ('p53', 'ON')), (('p53', 'ON'), ('p63', 'ON')), (('p53', 'ON'), ('p73', 'ON')), (('p53', 'ON'), ('miR203', 'ON')), (('p53', 'ON'), ('miR34', 'ON')), (('p53', 'ON'), ('miR200', 'ON')), (('p53', 'ON'), ('TGFbetaR', 'ON')), (('p53', 'ON'), ('AKT1', 'OFF')), (('p53', 'ON'), ('CTNNB1', 'OFF')), (('p53', 'ON'), ('DKK1', 'OFF')), (('p53', 'ON'), ('AKT2', 'OFF')), (('p53', 'ON'), ('RAC1', 'OFF')), (('p53', 'ON'), ('YAP1', 'OFF')), (('p53', 'ON'), ('PIK3CA', 'OFF')), (('p53', 'ON'), ('ERK', 'OFF')), (('p53', 'ON'), ('p21', 'OFF')), (('p53', 'ON'), ('SMAD', 'OFF')), (('p53', 'ON'), ('VIM', 'OFF')), (('p53', 'ON'), ('p63', 'OFF')), (('p53', 'ON'), ('p73', 'OFF')), (('p53', 'ON'), ('miR203', 'OFF')), (('p53', 'ON'), ('miR34', 'OFF')), (('p53', 'ON'), ('miR200', 'OFF')), (('p53', 'ON'), ('TGFbetaR', 'OFF')), (('p53', 'ON'), ('FAK', 'OFF')), (('p63', 'ON'), ('FAK', 'OFF')), (('p73', 'ON'), ('FAK', 'OFF')), (('miR34', 'ON'), ('FAK', 'OFF')), (('miR200', 'ON'), ('FAK', 'OFF'))])
```

In [75]:

```
filtered_res1 = filter_sensititivy(res_double, node="Cell_growth", maximum=0)
print(filtered_res.keys())
```

```
dict_keys([(('AKT1', 'ON'), ('p53', 'ON')), (('DKK1', 'ON'), ('p53', 'ON')), (('AKT2', 'ON'), ('p53', 'ON')), (('RAC1', 'ON'), ('p53', 'ON')), (('YAP1', 'ON'), ('p53', 'ON')), (('PIK3CA', 'ON'), ('p53', 'ON')), (('ERK', 'ON'), ('p53', 'ON')), (('p21', 'ON'), ('p53', 'ON')), (('SMAD', 'ON'), ('p53', 'ON')), (('VIM', 'ON'), ('p53', 'ON')), (('p53', 'ON'), ('p63', 'ON')), (('p53', 'ON'), ('p73', 'ON')), (('p53', 'ON'), ('miR203', 'ON')), (('p53', 'ON'), ('miR34', 'ON')), (('p53', 'ON'), ('miR200', 'ON')), (('p53', 'ON'), ('TGFbetaR', 'ON')), (('p53', 'ON'), ('AKT1', 'OFF')), (('p53', 'ON'), ('CTNNB1', 'OFF')), (('p53', 'ON'), ('DKK1', 'OFF')), (('p53', 'ON'), ('AKT2', 'OFF')), (('p53', 'ON'), ('RAC1', 'OFF')), (('p53', 'ON'), ('YAP1', 'OFF')), (('p53', 'ON'), ('PIK3CA', 'OFF')), (('p53', 'ON'), ('ERK', 'OFF')), (('p53', 'ON'), ('p21', 'OFF')), (('p53', 'ON'), ('SMAD', 'OFF')), (('p53', 'ON'), ('VIM', 'OFF')), (('p53', 'ON'), ('p63', 'OFF')), (('p53', 'ON'), ('p73', 'OFF')), (('p53', 'ON'), ('miR203', 'OFF')), (('p53', 'ON'), ('miR34', 'OFF')), (('p53', 'ON'), ('miR200', 'OFF')), (('p53', 'ON'), ('TGFbetaR', 'OFF')), (('p53', 'ON'), ('FAK', 'OFF')), (('p63', 'ON'), ('FAK', 'OFF')), (('p73', 'ON'), ('FAK', 'OFF')), (('miR34', 'ON'), ('FAK', 'OFF')), (('miR200', 'ON'), ('FAK', 'OFF'))])
```

In [76]:

```
for elem in filtered_res:
    for elem2 in filtered_res1:
        for elem3 in filtered_res2:
            if elem == elem2 and elem2 == elem3:
                print(elem)
```

```
(('AKT1', 'ON'), ('p53', 'ON'))
(('p21', 'ON'), ('p53', 'ON'))
(('p53', 'ON'), ('ERK', 'OFF'))
```

Conclusion: among the possible double mutation, the ones shown in the previous cell, can block both the the differentiation of epithelial cells into mesenchymal, both inhibits the growth of the tumor. Thus, these are possible candidates to simulate on PhysiboSS, to check if they effectively stop the tumor development.

In [ ]:
