## Supplementary File 3 for "Multiscale model of the different modes of cancer cell invasion"

variable\_analysis

This notebook is a companion for the article on a model describing the different modes of invasion using the tool PhysiBoSS.

PhysiBoSS is a modeling tool combining an agent-based approach for intercellular interactions and a Boolean model for the intracellular description.

The different parameters that are defined for the model show a variety of behaviors. By monitoring the values of some of the model variables, we show how the modes of invasion occur and evolve.

First, we define the libraries that are needed to perform the variable analyses

In [1]:

```
import matplotlib
import matplotlib.pyplot as plt
import math
import numpy as np
from matplotlib.widgets import Slider, Button
from ipywidgets import *
%matplotlib notebook
```

### Analysis of the evolution of the variables of the model¶

All the functions that allow cell interactions with the ECM, adhesion and evolution of the variables, have been written for this model ad hoc.

We focus on our analyses on a subset of parameters that regulate the main cell behaviors: adhesion, motility and degradation of the ECM. Their values range between [O, 1], since they all represent a percentage of activation:

- pintegrin: this parameter represents the percentage of integrins on the cell surface. The higher this value is, the more ECM the cell is able to degrade.
- pmotility: this parameter represents the tendency for a cell to move towards a certain direction. The higher this value is, the bigger the steps of the cell during migration will be.
- padhesion: this parameter represents the percentage of cell junction located on the surface of the cell membrane. This value can also be seen as a percentage of the amount of E-Cadherin present. In the modeling framework, depending on this value and the threshold selected by the user, the cells will develop more or less hook-like junctions and this effect will be linked to the possibility to form clusters of cells.

These variables evolve in time and are function of the state of the surrounding. They are based on the following equation:

$X\_{t+1} = X\_{t} + X\_{t}(1 - X\_{t})\frac{d\_{t}}{10} $

Where $(X\_{t})$ represents the value of the coefficient at step t, $X\_{t+1}$ at step t+1, and $d\_{t}$ the phenotype step.

#### An example for the phenotype ECM\_adhesion and the variable pintegrin¶

The phenotypes of individual cells will be modified according to these variables. As example, let us consider the read-out "ECM\_adhesion".

At each step of the simulation, the value of this output node is checked and the equation for the evolution of the associated variable is called (in this case the variable is "pintegrin"). The value of the phentoype is used to set the sign of $d\_{t}$.
If the value of the node is 1 (activation), $d\_{t}$ will have positive sign, if the value of the node is 0, $d\_{t}$ will have negative sign.
According to the sign, the curve associated to the equation changes its concavity, increasing or decreasing the value of the variable.

Below we report the evolution of the curve for both situations.

##### Equation with negative $d\_{t}$.¶

In [2]:

```
# Initial values are set for a negative dt

init_x = 1
dt = -6
time_step = np.arange(0, 11)

def update_param(x, time_step, dt):
    
    values = []

    for i in time_step:
    
        x += x*(1-x)*(dt/10)
    
        values.append(x)
        
    return values
```

The slider sets the initial condition of the variable

In [3]:

```
fig = plt.figure(figsize=(9, 6), dpi=100)
ax = fig.add_subplot(1, 1, 1)
line, = ax.plot(time_step, update_param(init_x, time_step, dt))
x_label = plt.xlabel("time_step")
y_label = plt.ylabel("parameter")
plt.xlim(0, 10)
plt.ylim(0, 1.1)


def update(w = 0.0):
    line.set_ydata(update_param(w , time_step, dt))
    fig.canvas.draw_idle()

interact(update, w=(0, 1, 0.01));
```

##### Equation with positive $d\_{t}$.¶

The slider sets the initial condition of the variable

In [4]:

```
init_x = 1
dt = 6
time_step = np.arange(0, 11)

fig = plt.figure(figsize=(9, 6), dpi=100)
ax = fig.add_subplot(1, 1, 1)
line, = ax.plot(time_step, update_param(init_x, time_step, dt))
x_label = plt.xlabel("time_step")
y_label = plt.ylabel("parameter")
plt.xlim(0, 10)
plt.ylim(0, 1.1)


def update(w = 0.0):
    line.set_ydata(update_param(w , time_step, dt))
    fig.canvas.draw_idle()

interact(update, w=(0, 1, 0.01));
```

As it can be verified, using the slider in the previous graph, both the curves have two singularities, the first one when $X\_{t+1} = 1$ and the second one when $X\_{t+1} = 0$. It is possible to verify this by setting the slider on 0 or 1.
In both conditions, the curve is saturated, which means that the variable has reached its asymptotic solution.

To solve this problem, we added a $\epsilon$ term, where $\epsilon$ << 1, that avoids saturation.

In [5]:

```
x = 0.99
dt = -6
time_step = np.arange(1, 20)

values = []

for i in time_step:
    
    x += x*(1-x)*(dt/10)
    
    values.append(x)
    
fig = plt.figure(figsize = (9, 5))

x_label = plt.xlabel("time_step")
y_label = plt.ylabel("parameter")

plt.plot(time_step, values)
```

Out[5]:

```
[<matplotlib.lines.Line2D at 0x7f27e74358e0>]
```

In [6]:

```
x = 0.01

dt = 6

time_step = np.arange(1, 20)

values = []

for i in time_step:
    
    x += x*(1.1-x)*(dt/10)
    
    values.append(x)
    
fig = plt.figure(figsize = (9, 5))

x_label = plt.xlabel("time_step")
y_label = plt.ylabel("parameter")

plt.plot(time_step, values)
```

Out[6]:

```
[<matplotlib.lines.Line2D at 0x7f27e7418340>]
```

#### Conclusion¶

The same scheme is applied to each of the model's variables described previously. The definition of these variables will be improved in the future and adapted to experimental observations.
In the future will be also possible to use a different function to describe the evolution of each variable, such as different Hill or Heaviside functions.

In [ ]:
